# Inhibitor of Differentiation 4 (ID4) represses myoepithelial differentiation of mammary stem cells through its interaction with HEB

**DOI:** 10.1101/2020.04.06.026963

**Authors:** Holly Holliday, Daniel Roden, Simon Junankar, Sunny Z. Wu, Laura A. Baker, Christoph Krisp, Chia-Ling Chan, Andrea McFarland, Joanna N. Skhinas, Thomas R. Cox, Bhupinder Pal, Nicholas Huntington, Christopher J. Ormandy, Jason S. Carroll, Jane Visvader, Mark P. Molloy, Alexander Swarbrick

## Abstract

Differentiation of stem cells embedded within the mammary epithelium is orchestrated by lineage-specifying transcription factors. Unlike the well-defined luminal hierarchy, dissection of the basal lineage has been hindered by a lack of specific markers. Inhibitor of Differentiation 4 (ID4) is a basally-restricted helix-loop-helix (HLH) transcription factor essential for mammary development. Here we show that ID4 is highly expressed in basal stem cells and decreases during myoepithelial differentiation. By integrating transcriptomic, proteomic, and ChIP-sequencing data, we reveal that ID4 is required to suppress myoepithelial gene expression and cell fate. We identify the bHLH protein HEB as a direct binding partner of ID4, and describe a previously-unknown role for this regulator in mammary development. HEB binds to E-boxes in regulatory elements of developmental genes, negatively regulated by ID4, involved in extracellular matrix synthesis and cytoskeletal contraction. Together our findings support a model whereby ID4 binds to HEB and blocks it from promoting myoepithelial specialisation. These new insights expand our current understanding into control of myoepithelial differentiation and mammary gland morphogenesis.

## Introduction

The mammary gland undergoes marked tissue remodelling throughout life. During murine pubertal development, terminal end buds (TEBS) located at the tips of the ducts invade into the surrounding stromal fat pad (Macias & Hinck, 2012, Williams & Daniel, 1983). This process is driven by collective migration and rapid proliferation of outer cap cells, which surround multiple layers of inner body cells (Williams & Daniel, 1983). As the ducts elongate, the cap cells differentiate into the basal cell layer and the body cells adjacent to the basal cells give rise to the luminal cell layer (Williams & Daniel, 1983), while the innermost body cells undergo apoptosis to sculpt the bilayered ductal tree present in the adult gland (Paine & Lewis, 2017). During pregnancy the gland undergoes alveolar morphogenesis in preparation for production and secretion of milk at lactation. The luminal cells differentiate into milk-producing alveolar cells and the milk is ejected from the gland by the contractile action of specialised myoepithelial cells, smooth muscle-like epithelial cells which differentiate from basal cells (Macias & Hinck, 2012). From hereon ‘basal’ refers to the basal lineage, encompassing cap cells, duct basal cells, and myoepithelial cells.

Initial transplantation experiments demonstrated that the basal lineage contains bipotent mammary stem cells (MaSCs), capable of forming both myoepithelial and luminal populations, while luminal cells are restricted in their differentiation potential (Shackleton, Vaillant et al., 2006, Stingl, Eirew et al., 2006). Subsequent lineage tracing experiments support the existence of a small population of basally-restricted bipotent stem cells (Rios, Fu et al., 2014, Wang, Cai et al., 2015), with a majority of unipotent basal and luminal mammary stem cells (Davis, Lloyd-Lewis et al., 2016, Lilja, Rodilla et al., 2018, Lloyd-Lewis, Davis et al., 2018, Prater, Petit et al., 2014, Scheele, Hannezo et al., 2017, van Amerongen, Bowman et al., 2012, Van Keymeulen, Rocha et al., 2011, Wuidart, Ousset et al., 2016, Wuidart, Sifrim et al., 2018) that replenish the myoepithelial and luminal lineages throughout life.

Lineage specifying transcription factors are responsible for directing luminal and myoepithelial differentiation, and also for maintaining the self-renewal of uncommitted stem cells upstream in the mammary epithelial hierarchy. Transcriptomic profiling of sorted epithelial subpopulations has been instrumental in identifying such lineage specifying transcription factors that regulate each step of luminal-alveolar differentiation (Asselin-Labat, Sutherland et al., 2007, Bouras, Pal et al., 2008, Buchwalter, Hickey et al., 2013, Carr, Kiefer et al., 2012, Chakrabarti, Wei et al., 2012, Kouros-Mehr, Slorach et al., 2006, Liu, Ginestier et al., 2008, Oakes, Naylor et al., 2008, Yamaji, Na et al., 2009). However, due to lack of specific cell markers that can resolve stem and myoepithelial populations, it has been challenging to dissect molecular regulators of basal differentiation. While a number of basal-specific transcription factors have been identified, such as P63, SLUG, SOX9, SRF and MRTFA (Guo, Keckesova et al., 2012, Li, Chang et al., 2006, Mills, Zheng et al., 1999, Sun, Boyd et al., 2006, Yang, Schweitzer et al., 1999), their role in the basal compartment and myoepithelial specialisation is poorly understood. ID proteins (ID1-4) are helix-loop-helix (HLH) transcriptional regulators that lack a DNA binding domain. They function by dimerizing with basic HLH (bHLH) transcription factors and preventing them from binding to E-box DNA motifs and regulating transcription (Benezra, Davis et al., 1990). E-box motifs are found in regulatory regions of genes involved in lineage specification and as such ID proteins and bHLH transcription factors are critical regulators of stemness and differentiation across diverse cellular linages (Massari & Murre, 2000). The expression of ID4 in mouse and human mammary epithelium is exclusive to the basal population (Lim, Wu et al., 2010). We and others have demonstrated that ID4 is a key regulator of mammary stem cells, required for ductal elongation during puberty (Best, Hutt et al., 2014, Dong, Huang et al., 2011, Junankar, Baker et al., 2015). ID4 was also shown to have a role in blocking luminal differentiation (Best et al., 2014, Junankar et al., 2015).The precise molecular mechanism by which ID4 functions in the mammary gland, including its full repertoire of transcriptional targets and interacting partners, have yet to be determined. Here we show that ID4 marks immature basal cells and demonstrate that ID4 inhibits myoepithelial differentiation. Moreover, using unbiased interaction proteomics, we identify a novel player in the mammary differentiation hierarchy – the bHLH transcription factor HEB. By mapping the genome-wide binding sites of HEB we show that it directly binds to regulatory elements of ID4 target genes involved in myoepithelial functions such as contraction and extracellular matrix (ECM) synthesis.

## Results

### Loss of ID4 causes upregulation of myoepithelial genes in basal cells

In order to determine the genes regulated by ID4 in mammary epithelial cells, we FACS-enriched basal, luminal progenitor and mature luminal cells from wild type (WT) and ID4 knockout (KO) mice (Yun, Mantani et al., 2004) and performed RNA-sequencing (RNA-seq) (Fig. 1a). No significant differences were observed in the proportion of mammary epithelial subpopulations between WT and KO mice (Fig. S1a). 96 genes were significantly (FDR<0.05) upregulated and 141 genes downregulated in ID4 KO basal cells. In contrast, only 2 genes were differentially expressed in the two luminal subpopulations (Fig. 1b-c and Table S1) suggesting that ID4 predominantly regulates gene expression within basal cells *in vivo*.

**Figure 1.**
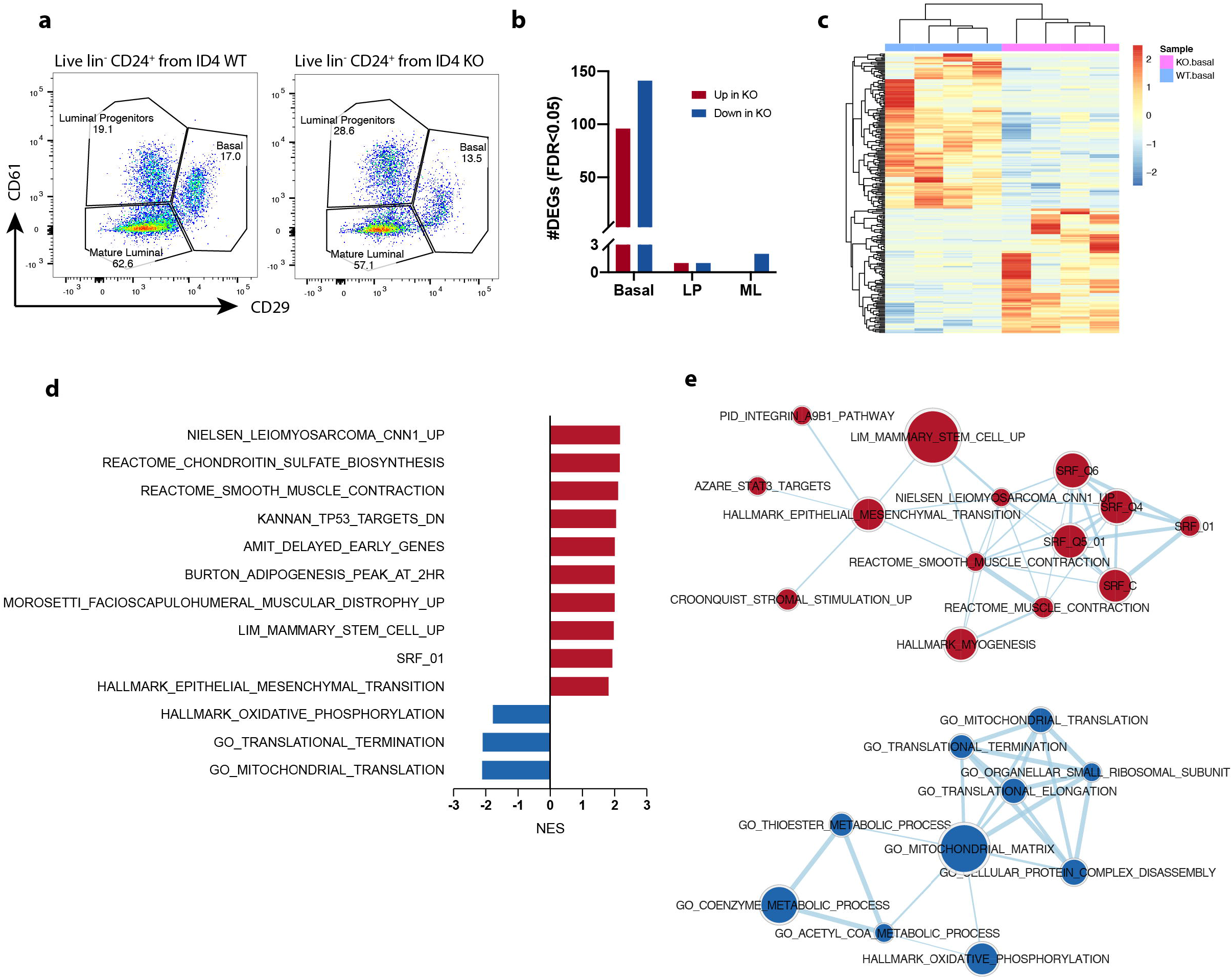
Loss of ID4 results in upregulation of myoepithelial genes and downregulation of cell growth genes in sorted mammary basal cells. **a)** Live lineage negative CD24^+^CD29^hi^CD61^+^ basal, CD24^+^CD29^lo^CD61^+^ luminal progenitor and CD24^+^CD29^lo^CD61^-^ mature luminal cells were isolated by FACS from adult (10-12 weeks) ID4 wild type (WT) and knockout (KO) mice at estrus for RNA-seq. Representative FACS plots shown from 4 experiments. **b)** Number of significantly (FDR<0.05) differentially expressed genes (DEGs) upregulated (red) or downregulated (blue) in ID4 KO epithelial subpopulations compared to ID4 WT. LP = luminal progenitor, ML = mature luminal. **c)** Heat map displaying the 237 significant differentially expressed genes between ID4 WT and ID4 KO basal cells. **d)** Genes were ranked based on the limma t-statistic comparing ID4 WT and KO basal cells and GSEA was carried out using the C2all, C3TF, C5 and Hallmark gene sets. The top 10 positive and negatively enriched pathways with an FDR<0.1 are displayed. Only 3 pathways were negatively enriched with an FDR<0.1. **e)** GSEA results were visualised using the EnrichmentMap plugin on Cytoscape. Nodes represent gene sets and edges represent overlap. Red nodes are positively enriched and blue nodes are negatively enriched. Gene sets with an FDR<0.25 are shown.

In line with the known role of ID4 in promoting proliferation of mammary epithelial cells (Dong et al., 2011, Junankar et al., 2015), the genes downregulated in ID4 KO basal cells were enriched for pathways involved in cell growth such as translation and metabolism (Fig. 1d-e and Fig. S1b). Conversely, the genes upregulated in KO basal cells were enriched for pathways related to basal cells and smooth muscle function. These included a muscle-derived leiomyosarcoma signature (Nielsen, West et al., 2002), smooth muscle contraction genes, epithelial-mesenchymal transition (EMT) and Serum Response Factor (SRF) targets (Fig. 1d and Fig. S1c), which shared common genes as indicated by the connecting edges in the enrichment map network (Fig. 1e). SRF target genes are of relevance as SRF is a master regulator of cytoskeletal contraction, and is one of the only transcription factors implicated in myoepithelial differentiation (Li et al., 2006, Miano, Long et al., 2007, Sun et al., 2006). Taken together, loss of ID4 causes basal cells to decrease proliferation and adopt a more differentiated myoepithelial and mesenchymal gene expression program, implicating ID4 in the repression of basal cell specialisation.

### ID4 marks basal stem cells and is dynamically regulated during myoepithelial differentiation

We previously showed that ID4 is heterogeneously expressed in basal cells and that ID4-positive cells have enhanced mammary reconstitution activity (Junankar et al., 2015). To further characterise the phenotype of ID4-positive basal cells, we used an ID4-GFP reporter mouse in which the ID4 promoter drives GFP expression (Best et al., 2014). Basal cells with high expression of the stem cell marker CD49f (also known as ITGA6) and the epithelial marker EPCAM have been shown to be enriched for MaSC activity (Prater et al., 2014, Stingl et al., 2006). ID4-GFP expression was maximal in this EPCAM^hi^ CD49f^hi^ subset (Fig. 2a). ID4-GFP expression within the basal gate was binned into 3 groups: bright (top 10%), intermediate (middle 80%) and dim (bottom 10%) and the median fluorescent intensity (MFI) of EPCAM and CD49f was analysed within these groups (Fig. 2b-c). ID4-bright cells had significantly higher EPCAM and CD49f MFI than ID4-dim and intermediate cells, suggesting ID4 marks basal stem cells.

**Figure 2.**
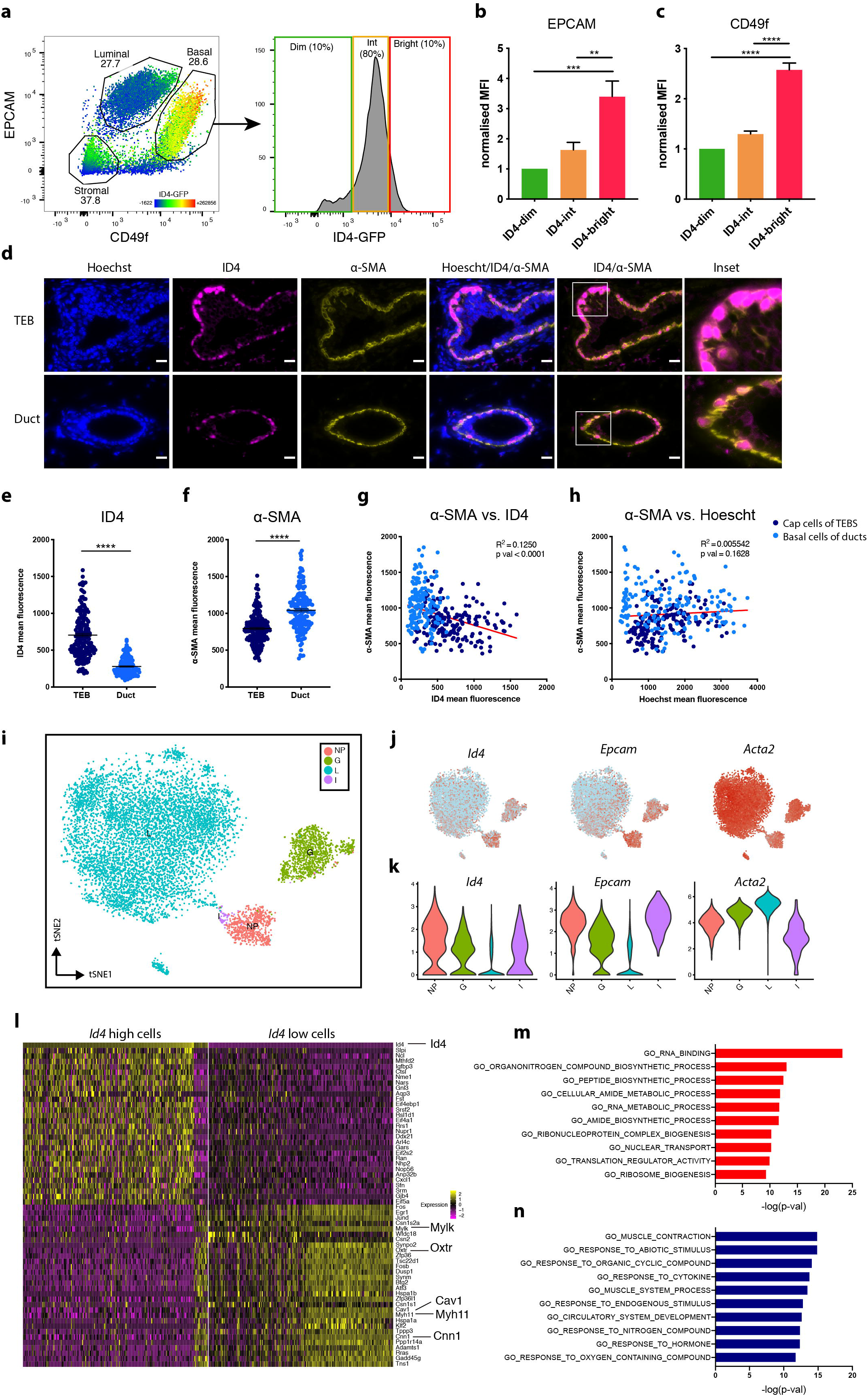
ID4 marks basal stem cells and decreases in terminally differentiated myoepithelial cells. **a)** FACS analysis of EPCAM and CD49f in live lin-mammary cells from adult (10-14 week) Id4floxGFP reporter mice. ID4-GFP expression is indicated in the heatmap scale. Representative plots from 5 experiments shown. Basal cells were binned into 3 groups based on ID4-GFP expression, ID4-bright (red), ID4-intermediate (orange) and ID4-dim (green) and the median fluorescence intensity (MFI) of EPCAM **(b)** and CD49f **(c)** were compared between the 3 gates. MFI expressed as a fold change relative to the ID4-low basal cells. Ordinary one-way ANOVA test was used to test significance. n=5. Error bars represent SEM. ** p<0.01, *** p<0.001, **** p<0.0001. **d)** Representative co-immunofluorescent staining of ID4 and α-SMA in TEB and duct from a pubertal (6 week) mammary gland. Scale bar = 20 μm. Comparison of ID4 **(e)** and α-SMA **(f)** mean fluorescence between individual cap cells (dark blue) and basal duct cells (light blue). Unpaired two-tailed students t-test. **** p<0.0001. Correlation between α-SMA and ID4 **(g)** and Hoescht **(h)** mean fluorescence in individual cap cells (dark blue) and basal duct cells (light blue). R^2^ and p values are displayed. Data is pooled from 9 mice. Approximately 20 TEB cap cells and 20 ductal basal cells were analysed per mouse. **i)** tSNE plot of 9663 Krt5+/Krt14+ basal cells from (Bach et al., 2017). 2 mice were analysed per developmental stage. NP = Nulliparous (8 week), G= Gestation (Day 14.5), L = Lactation (Day 6), I = Involution (Day 11). Feature plots **(j)** and Violin plots **(k)** displaying expression of *Id4, Epcam* and *Acta2* in single cells in the different developmental stages. **l)** Heatmap displaying top and bottom 30 differentially expressed genes between the top and bottom 200 *Id4* high and *Id4* low basal cells across all stages. Top 10 GO terms enriched in the top 50 genes upregulated in *Id4* high **(m)** and low **(n)** basal cells.

We next sought to locate ID4-high and ID4-low populations in a tissue context and to further investigate the association between ID4 and markers of myoepithelial differentiation. During ductal elongation, the cap cells differentiate at the neck of the TEBs and mature into myoepithelial cells which form the outer basal layer of the ducts (Paine & Lewis, 2017). Mammary gland sections from pubertal mice, in which TEBs and ducts are both present, were stained for ID4 and the myoepithelial marker alpha-smooth muscle actin (α-SMA). ID4 expression was highest in the nuclei of cap cells at the extremity of the TEBs and expressed at lower levels in basal cells of ducts (Fig. 2d and Fig. S2). Conversely, α-SMA expression was higher in ductal basal cells and lower in cap cells. ID4-high cap cells had a compact cuboidal epithelial appearance compared to the more separated elongated morphology of the ID4-low duct cells (Fig. 2d and Fig. S2). Quantification of fluorescence demonstrated that ID4 was more highly expressed in cap cells of TEBs than in basal cells of ducts, while the opposite was true for α-SMA expression (Fig. 2e-f). A negative correlation between α-SMA and ID4 was observed, with clear separation between cap (dark blue Fig. 2g) and duct basal cells (light blue Fig. 2g). As a negative control, α-SMA fluorescence was compared with nuclear stain Hoechst and no correlation or separation based on region was observed (Fig. 2h). Thus, based on marker expression, morphology and spatial localization, ID4 expression is high in epithelial-like cap cells and is lower in myoepithelial cells.

Terminal differentiation of basal cells into contractile myoepithelial cells occurs during lactation. We interrogated a published single cell RNA-seq (scRNA-seq) dataset (Bach, Pensa et al., 2017) to examine *Id4* expression dynamics over postnatal murine mammary gland development. In this study, individual EPCAM+ mammary epithelial cells from four developmental stages: nulliparous (8 week), gestation (day 14.5), lactation (day 6), and involution (day 11) were captured and profiled. We limited our analysis to basal cell clusters (9,663 cells), defined by expression of both *Krt5* and *Krt14*. Basal cells broadly clustered by developmental time point (Fig. 2i). Increased differentiation of myoepithelial cells appears to proceed from nulliparous to gestation to lactation, associated with decreased expression of epithelial marker *Epcam* and increased expression of *Acta2* (encoding α-SMA) (Fig. 2j-k), consistent with gradual acquisition of a smooth muscle phenotype and loss of adherent epithelial features (Deugnier, Moiseyeva et al., 1995). Like *Epcam, Id4* expression was highest in basal cells from nulliparous mice, and decreased in basal cells of pregnant and lactating mice (Fig. 2j-k). This result was validated on the protein level by immunohistochemical staining for ID4 on mammary gland sections at different developmental time points (Fig. S2b). Using the Monocle 2 package (Trapnell, Cacchiarelli et al., 2014), we performed pseudotemporal ordering of all basal cells to form a myoepithelial differentiation trajectory (Fig. S2c). As expected, the nulliparous and involution cells clustered together in pseudotime space in the least differentiated part of the trajectory. Basal cells from gestating mice were dispersed between the nulliparous and lactation stages, while basal cells at lactation were the most differentiated. *Id4* expression decreased, while several myoepithelial markers (*Acta2, Cnn1, Mylk, Myh11* and *Oxtr*) increased over pseuodotime (Fig. S2d).

We next sought to identify transcriptional signatures associated with high and low *Id4* expressing cells in the mammary basal epithelium across all developmental time points (Fig 2l and Table S2). Basal cells with high *Id4* expression were enriched for genes involved in RNA binding, metabolic processes, translation and ribosome biogenesis (Fig. 2m). Conversely, genes enriched in the *Id4*-low basal cells were involved in muscle contraction and response to cytokine and hormone stimuli, and circulatory system development (Fig. 2n). Notably, myoepithelial genes such as *Oxtr*, encoding the oxytocin receptor, and contractile genes *Mylk, Cnn1, Cav1* and *Myh11*, were among the top differentially expressed genes in the cells with low *Id4* expression (Fig. 2l and Table S2). Thus, ID4 downregulation during pregnancy and lactation is associated with terminal differentiation into functionally mature contractile myoepithelial cells.

### Loss of ID4 results in myoepithelial differentiation of mammary organoids

To functionally validate the role of ID4 in suppressing the myoepithelial differentiation of basal cells we generated a primary basal cell organoid model (Fig. 3a). The organoid model system complements and expands upon the findings from the KO mouse, as the acute consequence of ID4 loss on basal cell phenotype can be determined. Furthermore, organoids are less complex cellular systems than tissue, thus cell-autonomous effects can be isolated more precisely. ID4-positive basal cells from mice in which exons 1 and 2 of *Id4* are floxed (*Id4*^fl/fl^) (Best et al., 2014) were isolated using cell sorting. To overcome culture-induced senescence, basal cells were conditionally reprogrammed into a proliferative stem/progenitor state using an established protocol utilizing irradiated 3T3 fibroblast feeders and Rho Kinase (ROCK) inhibition (Liu, Ory et al., 2012, Prater et al., 2014) (Fig. 3a). Conditionally reprogrammed basal cells adopted an epithelial cobblestone morphology and expressed high levels of ID4 (Fig. S3a-b). Cells cultured in the absence of ROCK inhibitor or feeders adopted a flattened differentiated/senescent cell morphology and had reduced ID4 expression (Fig. S3a-b). Reprogrammed cells maintained basal marker expression of P63 and KRT14 (Fig. S3c) and could be grown as 3D organoids with basal marker KRT14 on the outer cell layer and luminal marker KRT8 on the inner cells of the organoids (Fig. S3d).

**Figure 3.**
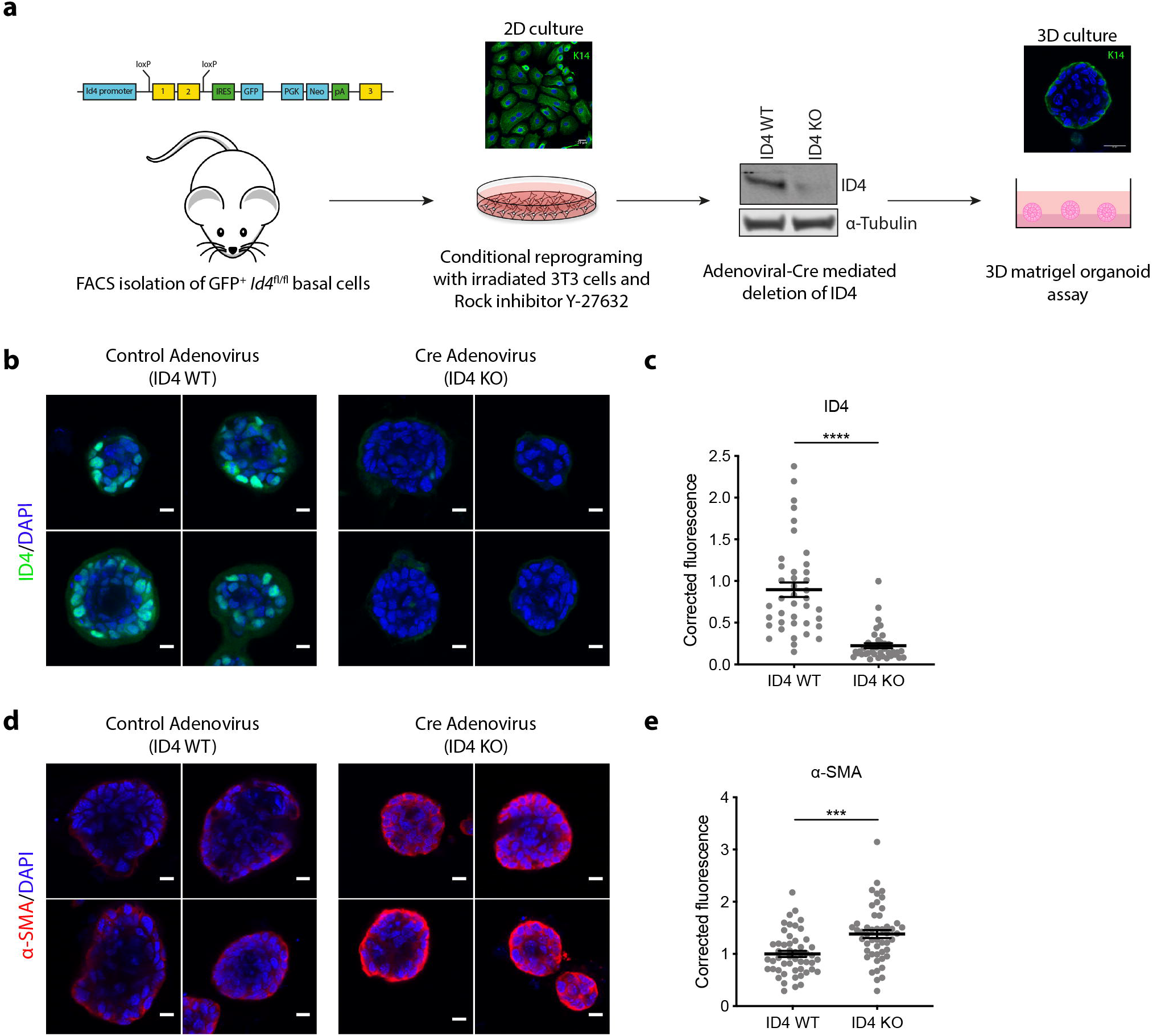
ID4 inhibits myoepithelial differentiation of organoids. **a)** Schematic diagram of 3D Matrigel organoid assay. ID4-GFP+ basal cells were FACS purified from adult (10-11 weeks) ID4-GFP reporter mice. Exon 1 and 2 of *Id4* are floxed and a GFP reporter cassette introduced. Basal cells are reprogrammed in culture using ROCK inhibitor Y-27632 and irradiated NIH-3T3 feeder cells. Adenoviral Cre is used to knock out ID4 as shown by western blotting. Single cells are then seeded on top of a Matrigel plug and grown for 6 days followed by immunofluorescent staining and quantification. Organoids grown from conditionally reprogrammed basal cells were treated with control GFP adenovirus (ID4 WT) or with Cre Adeno virus (ID4 KO) then stained for ID4 **(b)** and α-SMA **(d).** Scale bar = 10 μm. Fluorescence was quantified for ID4 **(c)** and α-SMA **(e)** in approximately 10 organoids per experiment. n=4. Unpaired two-tailed students t-test. *** p<0.001, **** p<0.0001.

ID4 expression was limited to the outer basal cells (Fig. 3b), recapitulating expression of ID4 in cap cells of TEBs (Fig. 2d). To test whether ID4 regulates differentiation of basal cells we deleted ID4 with Adenoviral-Cre and compared these to cells treated with control Adenoviral-GFP. ID4 protein was markedly downregulated in organoids infected with Cre adenovirus confirming successful gene deletion (Fig. 3b-c). Loss of ID4 resulted in increased α-SMA fluorescent intensity (Fig. 3d-e). This finding demonstrates that as well as marking undifferentiated basal cells, ID4 has a cell autonomous role in impeding maturation of basal cells into myoepithelial cells.

### ID4 inhibits expression of contractile and ECM genes *in vitro*

Given the negative association between ID4 and the differentiated myoepithelial phenotype, we next sought to determine direct ID4 target genes by performing RNA-seq following siRNA-mediated ID4 depletion in the spontaneously immortalized mouse mammary epithelial cell line Comma-Dβ (Danielson, Oborn et al., 1984). This normal-like cell line expresses basal markers and is commonly used as a model of mammary stem/progenitors cells as they retain the capacity of multi-lineage differentiation (Best et al., 2014, Danielson et al., 1984, Deugnier, Faraldo et al., 2006, Idoux-Gillet, Nassour et al., 2018, Junankar et al., 2015). They can also be readily expanded *in vitro* to produce a large amount of material for investigation of mechanisms by molecular biology and biochemistry. Western blotting confirmed 70-80% knockdown (KD) of ID4 protein 48 hr after siRNA transfection (Fig. 4a). RNA-seq analysis of ID4 KD cells resulted in 471 and 421 (FDR<0.05) down and upregulated genes, respectively, compared to the non-targeting siRNA control (Table S3).

**Figure 4.**
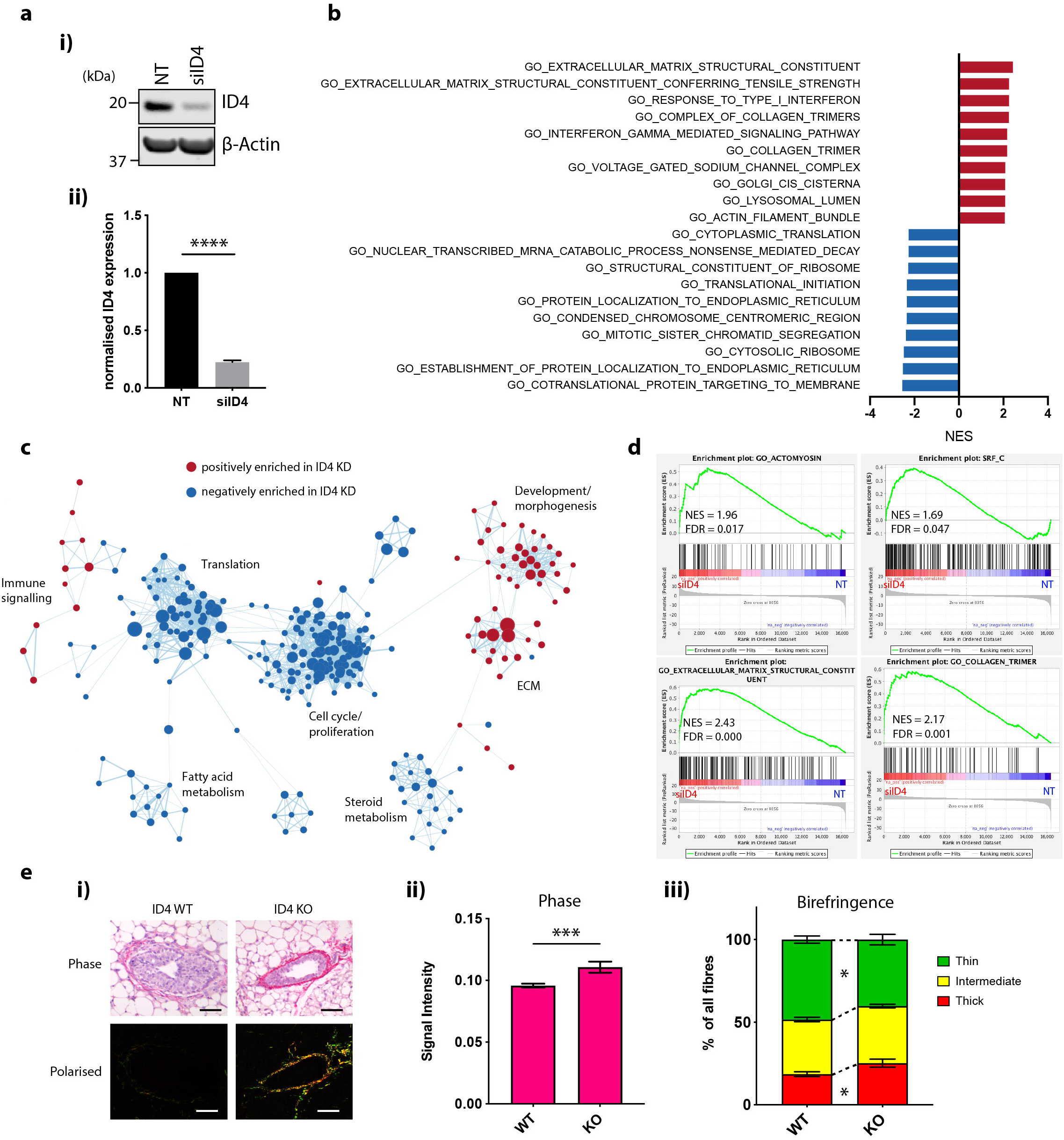
ID4 negatively regulates EMT and ECM production in mammary epithelial cells. **a)** Western blot analysis of ID4 expression in Comma-Dβ cells treated with non-targeting (NT) or ID4-targeting siRNA. Representative results from 3 western blots shown **(i).** Densitometry quantification of ID4 bands. Band intensity was normalised to β-Actin and expressed as fold change relative to NT control. N=3. Unpaired two-tailed students t-test. **** p<0.0001. Error bars represent SEM. **(ii). b)** Genes were ranked based on the limma t-statistic comparing NT and siID4 cells and GSEA was carried out using Gene Ontology (G) gene sets. The top 10 positively (red) and negatively (blue) enriched pathways are displayed **c)** GO GSEA results were visualised using the EnrichmentMap plugin on Cytoscape. Nodes represent gene sets and edges represent overlap. Red nodes are positively enriched and blue nodes are negatively enriched in siID4 cells compared to control NT cells. Gene sets with an FDR<0.1 are shown. **d)** Representative GSEA enrichment plots displaying the profile of the running Enrichment Score (green) and positions of gene set members on the rank ordered list. **e)** Collagen fibres were visualised by picrosirius red staining of TEBs from 6-week-old ID4 WT and KO mice. Scale bar = 50 μm (i). Total collagen staining **(ii)** and birefringence signal **(iii)** were quantified from approximately 3 TEBs from each mammary gland section. N=9 for WT and N=6 for KO mice. Unpaired two-tailed students t-test. * p<0.05. *** p<0.001. Error bars represent SEM.

Downregulated genes were predominantly involved in cell proliferation and growth pathways (Fig. 4b-c; blue) including hallmark gene sets such as E2F targets, MYC targets, and G2M checkpoint (Fig. S4a-b). Driving enrichment were several key cell cycle genes such as *Mki67, Cdk2, Cdk6* and *Cdk17* (Table S3). This result is consistent with the loss of cell growth gene expression programs in ID4 KO basal cells *in vivo* (Fig. 1d-e).

Genes acutely upregulated upon ID4 depletion were involved in development, morphogenesis, ECM remodelling, and immune signalling (Fig. 4b-c; red). Consistent with ID4 repressing myoepithelial specialisation, loss of ID4 resulted in upregulation of SRF targets, actomyosin cytoskeleton, EMT and myogenesis gene signatures (Fig. 4d and Fig. S4a-b). Several contractile genes were increased in ID4 depleted cells including *Cnn1, Cnn2, Tagln, Lmod1* and *Acta2* (FDR=0.06) (Table S3), many of which were inversely correlated with *Id4* expression in the scRNA-seq analysis (Fig. 2l and Table S2).

Several ECM genes encoding collagens (e.g. *Col1a1, Col1a2*, and *Col5a1)*, basement membrane laminins (e.g. *Lamc1* and *Lama4)*, and matricellular proteins (e.g. *Sparc)* were also upregulated upon ID4 depletion (Fig. 4c-d and Table S3). The ECM provides physical support to the mammary gland and is a source of biochemical signals important for coordinating morphogenesis (Muschler & Streuli, 2010). Changes in ECM gene expression are associated with EMT and cellular contractility (Kiemer, Takeuchi et al., 2001, Liu, Eischeid et al., 2012). To functionally validate the role of ID4 in regulating ECM proteins we examined whether ID4 represses ECM deposition *in vivo* using picrosirius red staining of mammary glands from ID4 WT and KO mice to visualise fibrillar collagen (Fig. 4e). ID4 KO TEBs in pubertal mammary glands were surrounded by a thickened collagen-dense ECM when compared to ID4 WT TEBs (Fig. 4e i-ii), indicating that ID4 normally restrains collagen expression by cap cells during puberty. In addition to total abundance, the thickness of bundled collagen fibres can be further distinguished using polarized light. Analysis of birefringence signal revealed an increase in thick fibres and a decrease in thin fibres in the ECM surrounding ID4 KO TEBs (Fig. 4e iii) signifying a redistribution of collagen composition, as well as an overall increase in collagen abundance, in the absence of ID4.

Hence, ID4 positively regulates proliferative genes and negatively regulates genes involved in myoepithelial functions such as contraction and ECM synthesis in mammary epithelial cells. These ID4-regulated functions are likely to be important for morphogenesis of the ductal tree during pubertal development.

### ID4 interacts with E-proteins in mammary epithelial cells

As ID4 lacks a DNA binding domain, it inevitably influences transcription through its interaction with other DNA-binding proteins. We used Rapid Immunoprecipitation Mass spectrometry of Endogenous proteins (RIME) to discover the binding partners of ID4 in Comma-Dβ cells (Mohammed, D’Santos et al., 2013). We identified 48 proteins that were significantly (p<0.05) more abundant in the ID4 IPs compared to the IgG negative control IPs in three independent RIME experiments (Fig. 5a and Table S4). ID4 was consistently identified in all replicates and was the top hit (Table S4), verifying the validity of the technique. Among the putative ID4 binding partners were many DNA and RNA interacting proteins (Fig. 5a).

**Figure 5.**
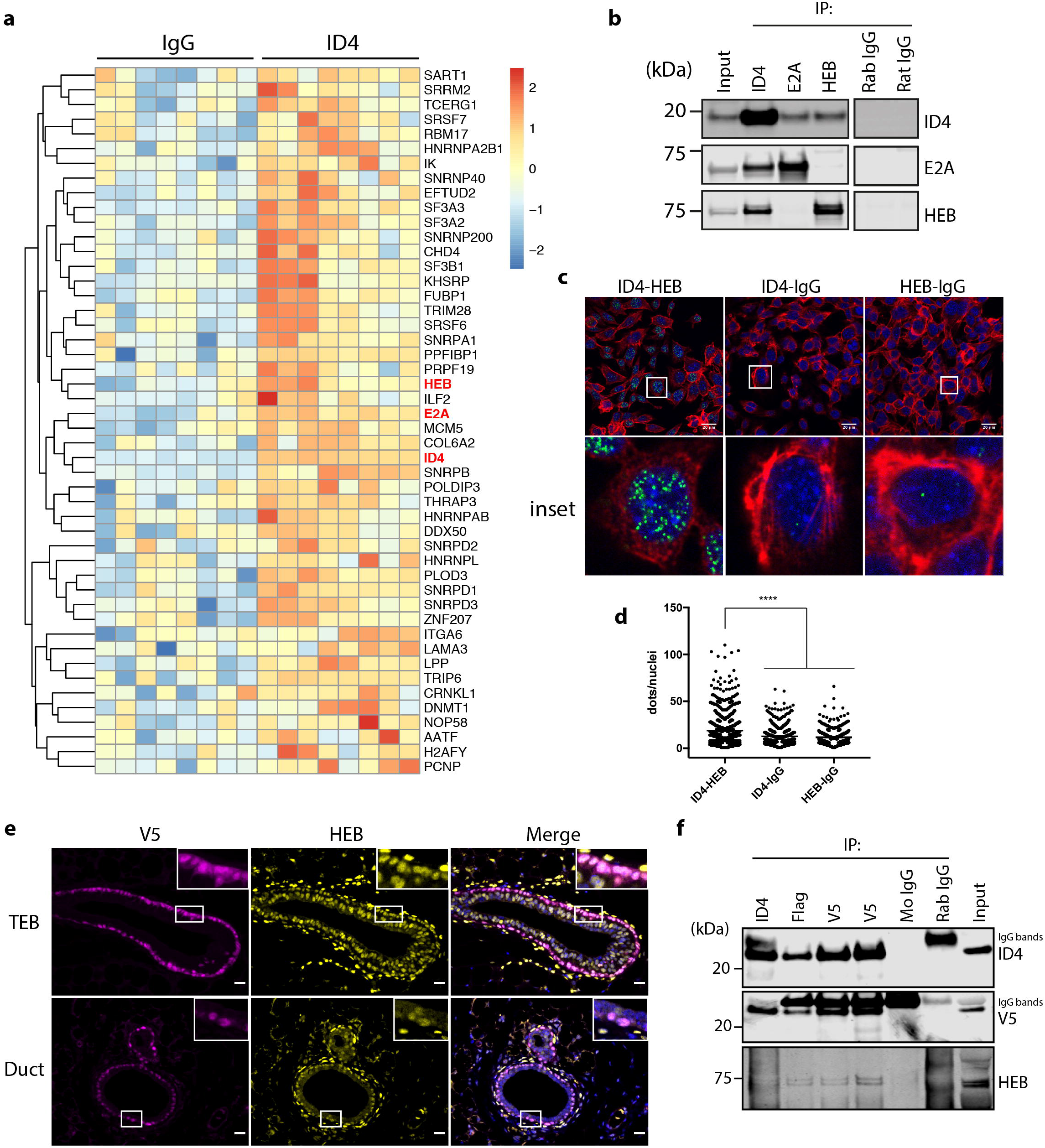
ID4 interacts with E-proteins E2A and HEB. **a)** Unsupervised hierarchical clustering heat map of SWATH RIME data from Comma-Dβ cells. Proteins with significantly higher abundance (p-value<0.05) in the ID4 IPs compared to IgG IPs are displayed. Log2 protein area was used to generate the heatmap. Data from 3 independent experiments shown, each with 2-3 technical replicates. **b)** Co-immunoprecipitation (co-IP) and western blotting of ID4, E2A and HEB and IgG negative controls from uncrosslinked Comma-Dβ lysates. **c)** Proximity ligation assay (PLA) in Comma-Dβ cells for ID4 and HEB or corresponding negative control IgG. The cytoskeleton is stained with phalloidin and nucleus with DAPI. Scale bar = 20 μm. High power insets are shown below. **d)** Quantification of PLA foci from 6 random fields of view for each condition, each with approximately 50 nuclei per image. Ordinary one-way ANOVA test was used to test significance. **** p<0.001. N=3. **e)** Co-immunofluorescent staining of V5 and HEB in TEB (upper) and duct (lower) from ID4-FlagV5 mouse mammary glands. High-power insets feature cells positive for both proteins. Scale bar = 20 μm. **f)** co-IP and western blotting from ID4-FlagV5 mammary gland protein extracts. ID4 was immunoprecipitated using antibodies raised against ID4 and Flag, and two independent V5 antibodies. ID4, V5 and HEB were detected by western blotting.

The E-proteins E2A and HEB were identified as ID4 interactors in our RIME analysis. There are three members of the E-protein family (E2A, HEB, and ITF-2), that dimerise with other E-proteins or tissue-specific bHLH transcription factors (e.g. MyoD and NeuroD) to regulate expression of lineage commitment genes (Wang & Baker, 2015). E2A has been implicated in branching morphogenesis of mammary organoids (Lee, Gjorevski et al., 2011), however, there are no previous studies implicating HEB in mammary gland development. Given this, and that E-proteins are the canonical binding partners of ID proteins in other lineages, we chose to pursue the ID4-HEB interaction further.

Reciprocal co-immunoprecipitation followed by western blotting (co-IP WB) experiments were performed to validate the ID4-E-protein interactions in the same Comma-Dβ cell line, as well as in normal human mammary epithelial cell lines PMC42 and MCF10A. IP of ID4 resulted in co-IP of E2A and HEB and correspondingly, IP of E2A and HEB resulted in co-IP of ID4 (Fig. 5b and S5a-b). E2A and HEB did not form heterodimers with each other in Comma-Dβ cells (Fig. 5b), implying that E-proteins either function as homodimers, or heterodimers with other bHLH proteins in this context. To identify mammary specific bHLH factors binding to HEB, we performed another RIME experiment by immunoprecipitating HEB protein. HEB (HTF4_MOUSE in Table S4) was identified as the top hit, and ID4 was also identified (Fig. S5c). However, we did not identify any other bHLH transcription factors in this experiment (Fig. S5c and Table S4). This suggests that HEB either binds DNA as a homodimer or binds another factor that was not detected by this assay. To independently validate the interaction between ID4 and HEB, the Proximity Ligation Assay (PLA) was used to visualize protein-protein interactions *in situ*. Multiple PLA foci were detected in the nuclei of Comma-Dβ cells co-stained with ID4 and HEB antibodies (Fig. 5c-d).

To test if ID4 and HEB interacted *in vivo, we* engineered a tagged ID4 mouse model in which a FlagV5 tag, a very efficient and specific target for immunoprecipitation, was integrated into the *Id4* locus downstream of the open reading frame. The result was FlagV5-tagged ID4 protein under the control of endogenous regulatory elements. We first ensured that ID4 and HEB were co-expressed by performing co-immunofluorescent staining for V5 and HEB. ID4-FlagV5 expression was tightly restricted to the cap and ductal basal cells, while HEB was more ubiquitous in its expression; detected in basal and luminal epithelial cells and surrounding stromal cells (Fig. 5e). ID4 and HEB co-localized in the nuclei of cap and duct basal cells (insets; Fig. 5e). Reciprocal co-IP was carried out from digested mammary glands of ID4-FlagV5 mice using antibodies raised against ID4, Flag, V5 (two different antibodies) and HEB (Fig. 5f). Each antibody tested was able to precipitate both ID4 and HEB, confirming interaction between the two transcription factors *in vivo*.

### HEB binds to regulatory elements of a subset of ID4 differentially expressed genes

To establish if ID4 regulates gene expression through HEB, we sought to determine if HEB directly binds to genes regulated by ID4 by performing ChIP-seq for HEB in Comma-Dβ cells. Across four biological replicates there were a total of 2752 HEB peaks identified (FDR<0.05). This was narrowed down to 956 consensus peaks which were present in at least 2 replicates. Transcription factor motif enrichment was carried out using MEME-ChIP (Machanick & Bailey, 2011) and the top enriched motifs were canonical E-box motifs (CANNTG), which are the binding sites for E-proteins (Fig. 6a). The majority of peaks were mapped to intergenic and intronic regions, and approximately 5% of the peaks occurred at gene promoters (Fig. S6a). We used the Genomics Regions Enrichment of Annotations Tool (GREAT) to analyse the functional significance of the regions bound by HEB (McLean, Bristor et al., 2010). In this unbiased analysis, top enriched pathways were related to actin cytoskeleton and ECM organization (Fig. 6b), resembling the pathways negatively regulated by ID4 in the gene expression profiling (Fig. 4b). This suggests that ID4 mediates repression though its physical interaction with HEB.

**Figure 6.**
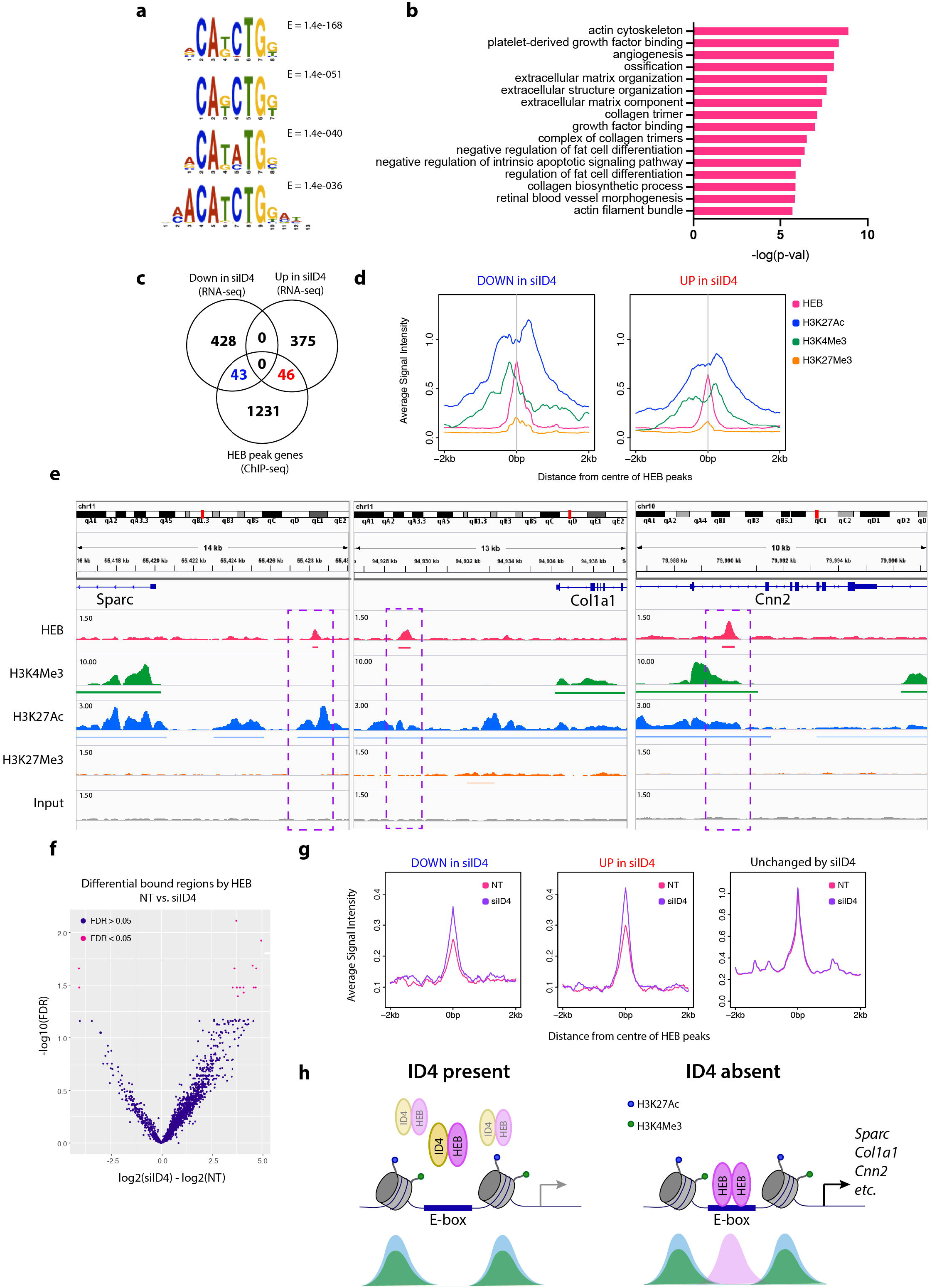
HEB directly binds to a subset of ID4 target genes. **a)** Top 4 enriched transcription factor binding motifs determined using MEME-ChIP for consensus HEB ChIP-seq peaks in Comma-Dβ cells. E-values are displayed. **b)** GREAT pathway analysis of consensus HEB peaks. Top 16 Gene Ontologies (Biological process, cellular component, molecular function) are displayed. **c)** Venn diagram showing overlap between genes associated with a HEB peak (GREAT basal plus extension annotation) and ID4 RNA-seq differentially expressed genes. **d)** Profile plots of average HEB, H3K4Me3, H3K27Ac, and H3K27Me3 signal intensity at regions associated with siID4 downregulated (left) and upregulated (right) RNA-seq differentially expressed genes. **e)** Examples of HEB and histone mark peaks occurring upstream of siID4 upregulated genes *Sparc, Col1a1* and *Cnn2* from the Integrative Genomics Viewer (IGV). Bars beneath peaks represent consensus MACS call (FDR<0.05) in at least 2 of 4 biological replicates. Input was used as a negative control. Purple boxes highlight HEB binding regions. Refseq genes shown in blue. Data scales for each track are indicated. **f)** Volcano plot of Differential Binding Analysis. Analysis using EdgeR of HEB binding in siID4 verses NT of 3 biological replicates. Regions with an FDR less than 0.05 are indicated in pink. **g)** Profile plots of average HEB signal intensity in NT control (pink) and siID4 (purple) conditions at regions associated with RNA-seq siID4 downregulated (left), upregulated (middle), and unchanged (right) genes. **h)** Model of ID4 and HEB action in mammary epithelial cells. Left: when ID4 is expressed it interacts with HEB, antagonising its transcriptional activity. Right: when ID4 is depleted, HEB dimerises and binds to E-box motifs in the promoters and enhancers of developmental genes involved in contraction and ECM. Below are hypothetical ChIP signals for H3K27Ac (blue), H3K4Me3 (green), and HEB (pink).

The consensus peaks were annotated to 1320 genes, using the default GREAT basal plus extension gene annotation rule (McLean et al., 2010). We overlapped these genes with those regulated following ID4 KD to determine the genes directly regulated by HEB (Fig. 6c). Approximately 10% of these genes had an associated HEB peak, which is more than expected by chance (p<5.65E-13; hypergeometric test). The remainder of the genes regulated by ID4 KD are likely due to secondary effects of ID4 KD or HEB independent mechanisms. The fact that there was a similar number of genes overlapping in both upregulated (46) and downregulated (43) genes, suggests that HEB can bind to sites both negatively and positively regulated by ID4. In line with this, E-proteins have previously been demonstrated to act as both activators and repressors of gene transcription by recruiting different co-factors (Bayly, Chuen et al., 2004, Zhang, Kalkum et al., 2004).

In parallel we performed ChIP-seq for three histone modifications; H3K4Me3 (active promoter mark), H3K27Ac (active enhancer mark) and H3K27Me3 (repressive chromatin mark) to elucidate the chromatin context of HEB-bound regions. A number of HEB peaks associated with ID4 regulated genes demonstrated a bimodal distribution of H3K27Ac signal (Fig. 6d and Fig. S6b) suggesting that HEB binds to enhancers. Some of the peaks were also localized to active promoters (Fig. 6d and Fig. S6b). HEB peaks were observed at enhancer-marked chromatin upstream of ID4-repressed ECM genes expressed in myoepithelial cells (Barsky & Karlin, 2005) such as *Sparc, Col1a1, Col1a2, Col3a1* and *Col5a1* (Fig. 6e and Fig. S6c). HEB binding was also observed near the promoter of contractile gene *Cnn2* (Fig. 6e). Of relevance *Cnn2*, encoding Calponin 2, was recently discovered to be regulated by a super enhancer specifically accessible in myoepithelial cells (Pervolarakis, Nguyen et al., 2019). This further indicates that ID4 inhibits HEB’s ability to activate transcription of genes that define the myoepithelial fate.

We next performed HEB ChIP-seq on Comma-Dβ cells in which ID4 had been depleted by siRNA in order to determine whether HEB DNA binding is augmented when released of inhibition by ID4. Western blotting revealed that ID4 protein was reduced to approximately 20% of control levels, while HEB expression was unchanged (Fig. S6d). E-box motifs were again enriched in both conditions (Fig. S6e). Differential binding analysis revealed a total of 290 regions changing upon ID4 depletion (p<0.05) (Table S5). More peaks were increased than decreased (263 compared to 27) suggesting that, as expected, depletion of ID4 increased HEB’s DNA binding activity (Fig. 6f and Fig. S6f). GREAT analysis revealed that the peaks that increased were involved in processes such as gland morphogenesis, skeletal development, and branching morphogenesis (Fig. S6g). No pathways were enriched in the regions that were decreased when ID4 was knocked down. Finally, we observed an increase in HEB binding in cells depleted of ID4, specifically at genes that were differentially expressed by ID4 KD (Fig. 6g).

Together, our ChIP-seq analysis suggests that HEB binds to E-box motifs in regulatory elements of basal developmental genes involved in ECM and the contractile cytoskeleton, and this is antagonised by its interaction with ID4 (Fig. 6h).

## Discussion

The mammary gland is the defining morphological feature of mammals and is also the source of one of the most common human cancers. Beyond their role in milk ejection at lactation, myoepithelial cells can also sense, synthesize and remodel the ECM, and control the polarity of neighbouring luminal cells (Barsky & Karlin, 2005, Gudjonsson, Rønnov-Jessen et al., 2002, Ingthorsson, Hilmarsdottir et al., 2015). Furthermore, myoepithelial cells are “natural tumour suppressors”; acting as physical barriers to prevent cancer cell invasion, secreting factors that limit growth and rarely undergoing malignant transformation (Gudjonsson, Adriance et al., 2005, Sternlicht, Kedeshian et al., 1997). Thus, understanding the regulation of the basal lineage has implications for fundamental mammary gland biology as well as breast cancer.

Both bipotent and unipotent stem cells have been identified within the mammary epithelium (reviewed in (Visvader & Clevers, 2016)). ID4 has previously been demonstrated to block luminal commitment of basal cells via inhibition of key luminal driver genes including *Elf5, Notch, Brca1, Esr1, PR* and *FoxA1* (Best et al., 2014, Junankar et al., 2015). We now show that ID4 can also block genes associated with myoepithelial differentiation, in part through the inhibition of the E-protein HEB. The dual inhibition of both luminal and myoepithelial differentiation by ID4 may protect the stem-cell phenotype of uncommitted basal cells during development. Subsequent downregulation of ID4, through a currently unknown mechanism, may then allow basal cells to adopt a luminal or myoepithelial fate depending on the cellular context.

HEB has been implicated in the specification of lymphocyte (Braunstein & Anderson, 2012), haematopoietic (Li, Brauer et al., 2017), mesodermal (Yoon, Foley et al., 2015), neuronal (Mesman & Smidt, 2017), and skeletal muscle (Conway, Pin et al., 2004) lineages. It however has not been associated with lineage commitment of epithelial tissues. Our proteogenomic analyses support the model outlined in Fig. 6h. When ID4 is highly expressed, such as in cap cells, it sequesters HEB off chromatin, preventing expression of differentiation genes, thus determining a stem-like state (Fig. 6h; left). When ID4 expression is low, HEB is able to bind to E-box DNA motifs at promoters/enhancers to activate transcription of developmental genes that specify functional myoepithelial cells (Fig. 6h; right). Further functional studies are needed to demonstrate HEB’s requirement in promoting myoepithelial differentiation.

Compared with the luminal lineage, the molecular regulators controlling the basal lineage remain poorly understood. One of the few transcription factors known to promote myoepithelial differentiation is MADS-box protein SRF and its associated co-activator MRTFA (Li et al., 2006, Sun et al., 2006). Interestingly, HEB and SRF cooperatively activate transcription of *Acta2* in cultured fibroblasts (Kumar, Hendrix et al., 2003). This occurred in an E-box dependent manner and was inhibited by ID1 and ID2 overexpression (Kumar et al., 2003). While we show that ID4 suppresses *Acta2, we* did not observe HEB binding to the *Acta2* promoter. However, it is possible that in the absence of ID4, HEB and SRF cooperate to drive expression of other myoepithelial genes. This is supported by the positive enrichment of SRF targets upon ID4 depletion. Subsequent studies should test whether HEB and SRF cooperate in mammary epithelial cells.

The newly elucidated roles of ID4 in regulating myoepithelial commitment and ECM deposition expand upon why ID4 is required for pubertal mammary gland morphogenesis (Best et al., 2014, Dong et al., 2011, Junankar et al., 2015). TEBs undergo collective migration, enabling the coordinated movement of adherent cells into the stromal fat pad (Ewald, Brenot et al., 2008). We hypothesise that the high levels of ID4 in cap cells prevent epithelial cells from acquiring a mesenchymal/myoepithelial phenotype. Similar mechanisms have been observed for the transcriptional repressor OVOL2, which inhibits EMT to allow for collective migration (Watanabe, Villarreal-Ponce et al., 2014), and for C/EBPα, which maintains epithelial homeostasis of human mammary epithelial cells (Lourenço, Roukens et al., 2020). During ductal elongation, collagenous stromal ECM is absent directly in front of the invading TEBs (Paine & Lewis, 2017, Silberstein, Strickland et al., 1990, Sternlicht, 2006). We show that ID4 suppresses collagen synthesis and deposition around TEBs, which may otherwise act as a physical barrier to impede invasion. In support of this idea, ectopic deposition of collagen by mammary epithelial cells induced by exogeneous TGF-β, or forced expression of recombinant type I collagen which is resistant to collagenase attack, causes ensheathment of TEBs by collagen, and retardation of ductal elongation (Feinberg, Zheng et al., 2018, Silberstein et al., 1990).

Developmental transcription factors are often dysregulated in cancer. ID4 is highly expressed in ~50% of basal-like breast cancer (BLBC) cases and associates with poor prognosis (Baker, Holliday et al., 2016, Junankar et al., 2015). Given the cell-intrinsic role of ID4 in promoting growth/proliferation and inhibiting differentiation in the mammary gland, it is easy to envision how overexpression of ID4 could lead to an aggressive breast cancer phenotype. Additionally, the suppression of ECM synthesis may allow tumour cells to easily invade into the surrounding stroma, akin to the invasion of cap cells during ductal elongation. Future work should test whether the mechanisms discovered here are conserved in cancer. If so, targeting ID4 may represent an attractive therapeutic strategy for ID4-positive BLBCs. Excitingly, a small-molecule Pan-ID inhibitor has recently been identified by the Benezra laboratory, which works by inhibiting the interaction between ID and E-proteins (Wojnarowicz, Lima E Silva et al., 2019).

To conclude, these new insights into ID4 and HEB function have helped unravel regulation within the elusive basal differentiation hierarchy, with broader implications for the regulation of epithelial stem cells in general, and also in tumour progression.

## Materials and methods

### Mice

All mice experiments were performed in accordance with the ethical regulations of the Garvan Institute Animal Experimentation Committee. ID4 KO mice were generated as previously described (Yun et al., 2004). The founder ID4-FlagV5 mouse was engineered using Crispr/Cas9 gene editing by the Mouse Engineering at Garvan/ABR (MEGA) facility. The ID4floxGFP mice were generated as previously described (Best et al., 2014). All mice used were on the FVB/N background. For the gene expression profiling experiment, mice were synchronised in estrus to reduce hormone-induced gene expression variation (Dalal, Estep et al., 2001), and checked by vaginal swab cytology.

### Mammary epithelial cell preparations

Mammary epithelial cells were prepared from freshly harvested 3^rd^ and 4^th^ mammary glands pooled from 4-8 female mice at the indicated ages. Glands were mechanically disrupted using a McIlwain tissue chopper then digested with 15,000 U collagenase (Sigma-Aldrich) and 500 U hyaluronidase (Sigma-Aldrich) in FV media (DMEM/F12 (Gibco), 5% (v/v) FBS (HyClone) 10 mM HEPES (Gibco), 0.14 IU/mL Insulin (Novo Nordisk), 500 ng/mL Hydrocortisone (Sigma-Aldrich), 20 ng/mL Cholera Toxin (Sigma-Aldrich), 2 ng/mL mEGF (Life Technologies)). Digestion was carried out in a shaking incubator for 1 hr at 37°C 225 rpm. The resulting organoids were further digested with warm 0.25% trypsin for 1 min with constant pipetting followed by treatment with 5 mg/mL Dispase (Roche) for 5 min in at 37°C. Cells were incubated with 1X red blood cell lysis buffer (BD Biosciences) for 5 min at room temperature. Cells were passed through a 70 μm strainer followed by a 40 μm strainer to remove clumps and debris. Live cells were counted using trypan blue (Thermo Fisher Scientific) and a haemocytometer prior to use in downstream applications.

### Flow cytometry and FACS

Single-cell suspensions of mouse mammary epithelial cells were blocked in Fc block cocktail (FACS buffer, 6.25 μg/mL Mouse BD Fc block (BD Bioscience) and 200 μg/mL Rat Gamma Globulin (Jackson ImmunoResearch)) for 10 min on ice. Fluorophore- or biotin-conjugated antibodies were diluted in Fc block cocktail and incubated on the cells on ice for 20 min in the dark. Antibodies used were CD24-PE (1:200, BD Biosciences, Clone M1/69), EPCAM-PerCP/Cy5.5 (1:200, Biolegend, Clone G8.8), CD29-APC/Cy7 (BioLegend, Clone HMβ1-1), CD61-APC (1:50, Thermo Fisher Scientific, Clone HMβ3-1), CD49f-APC (1:100, BioLegend, Clone GoH3), Ter119-biotin (1:80, BD Biosciences, Clone TER119), CD45-biotin (1:100, BD Biosciences, Clone 30-F11), CD31-biotin (1:40, Biolegend, Clone 390) and BP1-biotin (1:500, Thermo Fisher Scientific, Clone 6C3). Cells were washed and stained with streptavidin-BV421 (1:100, Biolegend). Following washing, cells were resuspended in FACS buffer at a density of 1×10^7^ cells/mL and DAPI was added (1:1000, Invitrogen). FACS was performed on a FACS Aria III 4 laser 15 colour sorter with BD FACS DIVA software. For analytical flow cytometry, a CytoFLEX 3 laser 13 colour flow cytometer (Beckman Coulter) was used. All flow cytometry data was analysed using FlowJo software version 10 (Tree Star Inc).

### RNA-seq

RNA was extracted using the Qiagen miRNease mini kit (Qiagen) for Comma-Dβ cells or the Qiagen miRNease micro kit (Qiagen) for FACS-sorted mammary epithelial cells. Cells were sorted directly into 700 μL Qiazol lysis reagent to minimise cell loss, with a maximum of 5×10^4^ cells per 1.5 mL Eppendorf tube. RNA was eluted in RNAse free water. The Qubit RNA BR Assay kit (Thermo Fisher Scientific) was used to measure RNA concentration and RNA integrity was determined using the Agilent Bioanalyser 2100 with the 6000 Nano Assay (Agilent Technologies).

For low input RNA extracted from FACS sorted mammary glands, the Ovation RNA-seq System V2 kit (NuGEN) was used to synthesise cDNA with RNA inputs ranging from 0.5-2 ng as per the manufacturers protocol. The Ovation Ultralow System V2 kit was then used to prepare libraries from the cDNA. cDNA libraries were sequenced on a HiSeq 2500 system (high output mode) (Illumina), with 125 bp paired-end reads. For RNA extracted from Comma-Dβ cells, The Illumina TruSeq Stranded mRNA Library Prep Kit (Illumina) was used with 1 μg of input RNA following the manufacturer’s instructions. cDNA libraries were sequenced on the NextSeq system (Illumina), with 75 bp paired-end reads.

Quality control was checked using FastQC (Andrews, 2010) to remove poor quality reads. Illumina sequencing adapters were then trimmed using Cutadapt (Martin, 2011). Reads were then aligned to the mouse reference genome mm10 using STAR ultrafast universal RNA-seq aligner (Dobin, Davis et al., 2013). RSEM accurate transcript quantification for RNA-seq data was used to generate a gene read count table and to filter genes with read counts of 0 (Li & Dewey, 2011). Differential gene expression analysis was performed using edgeR (McCarthy, Chen et al., 2012, Robinson, McCarthy et al., 2010) and voom/limma (Ritchie, Phipson et al., 2015) R packages. Genes were ranked based on the limma moderated t-statistic and this was used as input GSEA pre-ranked (Subramanian, Tamayo et al., 2005) using Molecular Signature Database (MSigDB) collections (v7.1). The EnrichmentMap Cytoscape plugin (Merico, Isserlin et al., 2010) was used to visualise the GSEA results.

### Analysis of public single-cell RNA-seq data

Raw Unique Molecular Identifier (UMI) expression data was taken from Bach *et al*. and re-processed using Seurat v2 using default parameters, as recommended by the developers (Satija, Farrell et al., 2015). All basal cell clusters, defined by *Krt5* and *Krt14* expression, were extracted for subsequent analysis. Developmental trajectories were inferred using Monocle 2 (Qiu, Mao et al., 2017). Clustering was first performed using the ‘densityPeak’ and ‘DDRTree’ methods, with a delta local distance threshold of 2, and a rho local density threshold of 3. For inferring pseudotime, we first selected all genes with a mean expression greater than 0.5, and an empirical dispersion greater than 1, and proceeded with the top 1000 significant genes for ordering. Differential gene expression between the two branch states were performed using the MAST algorithm through the Seurat package (Finak, McDavid et al., 2015, Satija et al., 2015). For comparisons between *Id4*-high and *Id4*-low states, we first filtered for cells with the detection of *Id4* to avoid technical biases from gene drop out. Differential gene expression was performed (as described above) between the top and bottom 200 cells grouped based on log normalised gene expression of *Id4*.

### Immunohistochemistry and immunofluorescence on mammary tissue sections

IHC staining was performed on 4 μm sections of formalin-fixed paraffin-embedded (FFPE) tissue. Slides were dewaxed with xylene and hydrated through graded alcohols. Antigen retrieval was performed for 1 min in a pressure cooker in DAKO target retrieval reagent s1699. A DAKO Autostainer was used for subsequent steps. Briefly, slides were incubated with DAKO peroxide block for 5 min and blocked with DAKO protein block for 30 min. ID4 Primary antibody (1:400, Biocheck BCH9/82-12) was incubated on the slides for 60 min. Following washing, slides were incubated with Envision rabbit secondary antibody (Agilent Technologies, Santa Clara, CA, USA) for 30 min. Slides were washed then incubated with DAKO DAB+ (Agilent Technologies) reagent for 10 min. Slides were then rinsed and counterstained with haematoxylin for 20-30 sec and dehydrated through graded alcohols, cleared using xylene and mounted using Ultramount #4 (Fronine). Bright-field images were captured using a Leica DM 4000 microscope with high-resolution colour camera (DFC450).

Immunofluorescence was performed manually on antigen retrieved tissue sections prepared as described for IHC. Slides were blocked with Mouse on Mouse (MOM) blocking buffer (Vector Biolabs) for 1 hour. Primary antibody was diluted in MOM diluent and incubated overnight at 4°C. Antibodies used were ID4 (1:400, Biocheck BCH9/82-12), α-SMA (1:200, Sigma-Aldrich A5228), HEB (1:200, Protein Tech 14419-1-AP), V5 (1:200, Santa Cruz sc-58052). Following washing, slides were incubated with fluorescent secondary antibody (1:500, Jackson ImmunoResearch) for 1 hr at room temperature. Nuclei were stained using Hoechst 33342 (Sigma-Aldrich). Slides were mounted using Prolong Diamond mounting media (Thermo Fisher Scientific). Fluorescent images were captured using a Leica DM 5500 microscope.

For quantification of fluorescence of ID4 and α-SMA in cap and ductal basal cells, 3 representative ducts and TEBs were imaged from each mouse (n=9). Using FIJI software, a region of interest was drawn around 10 randomly selected basal cells from each image. The mean fluorescence of ID4 and α-SMA was determined within each cell.

### Picrosirius red staining

Picrosirius red staining was performed following as per (Vennin, Chin et al., 2017, Vennin, Mélénec et al., 2019). Briefly, dewaxed FFPE sections were stained with haematoxylin. Sections were then treated with 0.2% Phosphomolybdic acid (Sigma-Aldrich) followed by 0.1% Sirius red (Sigma-Aldrich) Picric Acid-Saturated Solution (Sigma-Aldrich). Slides were then rinsed with acidified water (90 mM glacial acetic acid) then with 70% ethanol. A Leica DM 4000 microscope was used to image total collagen staining in the tissue sections by phase contrast. To image birefringence of collagen fibres, 2 polarised filters were used. Matched phase-contrast and polarised images were taken for each region of interest. Image analysis was performed using FIJI image analysis software.

### Proximity Ligation Assay

Comma-Dβ cells were grown on glass coverslips in 6 well plates until ~80% confluent. Cells were fixed in 4% Paraformaldehyde (PFA) (ProSciTech) for 15 min then permeablised in 0.2% Triton-X-100 for 15 min. The Duolink Proximity Ligation Assay (PLA) (Sigma-Aldrich) was performed as per the manufacturer’s protocol and using the following antibodies: ID4 (1:50, Santa Cruz sc-365656) and HEB (1:200, Protein Tech 14419-1-AP), and equivalent concentrations of IgG negative controls (sc-2027 and sc-2025). DAPI (Life Technologies) and Phalloidin (Life Technologies) were added to the final wash step. Coverslips were mounted onto glass slides with Prolong Diamond mountant (Thermo Fisher Scientific).

Cells were imaged using a Leica DMI Sp8 confocal microscope (63X oil objective). Six random fields of view per image were captured, each with approximately 50 cells, and images were quantified using Andy’s algorithms FIJI package (Law, Yin et al., 2017) to enumerate the number of PLA foci per nuclei.

### Cell lines

The mouse mammary epithelial cell line Comma-Dβ was a gift was a gift from Joseph Jeffery (University of Massachusetts, Amherst, MA, USA). Comma-Dβ cells were maintained in DMEM/F12 media (Gibco) supplemented with 2% FBS (HyClone), 10 mM HEPES (Gibco), 0.125 IU/mL Insulin (Novo Nordisk) and 5 ng/mL mEGF (Life Technologies). The human mammary epithelial cell line PMC42 was a gift from Professor Leigh Ackland (Deakin University, Melbourne, Victoria, Australia). PMC42 cells were maintained in RPMI 1640 (Gibco) supplemented with 10% FBS (HyClone). The MCF10A cell line was obtained from the American Type Culture Collection and were maintained in DMEM/F12 (Gibco), 5% Horse Serum (Thermo Fisher Scientific), 20 ng/mL hEGF (In Vitro Technologies), 0.5 mg/mL Hydrocortison (Sigma-Aldrich), 100 ng/mL Cholera Toxin (Sigma-Aldrich) and 0.125 IU/mL Insulin (Novo Nordisk).

### Conditional reprogramming of primary mouse basal cells

Viable basal cells were purified by FACS from 10-12 week old mice as described above. Cells were collected into FAD media (DMEM/F12 3:1 (Gibco) supplemented with 10% FBS, 0.18 mM Adenine (Sigma-Aldrich), 500 ng/mL Hydrocortisone (Sigma-Aldrich), 8.5 ng/mL Cholera Toxin (Sigma-Aldrich), 10 ng/mL mEGF (Life Technologies), 0.14 IU/mL Insulin (Novo Nordisk), 5 μM Y-27632 (Seleckchem), 1X AB/AM (Gibco) and 50 μg/mL Gentamicin (Life Technologies). Basal cells were maintained in culture using as per (Prater et al., 2014). Tissue culture flasks coated with Growth Factor-reduced Matrigel (Corning) diluted 1:60 in PBS for 30 min to 1 hr at 37°C. Excess Matrigel/PBS solution was aspirated from the flasks and basal cells were seeded at a density of ~5000 cells/cm^2^. Basal cells were co-cultured with irradiated (50 Gy) NIH-3T3 feeder cells at a density of 10,000 cells/cm^2^. Cells were maintained in a 37°C low 5% oxygen 5% CO_2_ incubator. Differential trypsinisation was used to first remove the less-adherent 3T3 cells when passaging.

### Organoid culture

A 40 μL plug of Matrigel (Corning) was added to each well of 8-well chamber slides (Corning) and allowed to set for 30 min in a 37°C incubator. Primary cells were resuspended in Epicult-B (Stemcell Technologies) media containing 2% Matrigel and 12,000 cells were seeded into each chamber in a total volume of 400 μL. Chambers were observed using a light microscope to ensure that cells were not aggregated. Organoids were allowed to form over 1 week and media was changed every 3 days.

### Immunofluorescent staining of organoids

Organoids were stained within the chamber slides. Media was aspirated and organoids were fixed with 2% PFA diluted in PBS for 20 min at room temperature. Organoids rinsed with PBS for 5 min following permeabilisation with 0.5% Triton X-100 in PBS for 10 min at 4°C then rinsed 3 times with 100 mM Glycine (Astral Scientific) in PBS for 10 min each. Organoids were blocked for 1 hr in IF buffer with 10% goat serum (Vector Laboratories). Primary antibody made up in blocking buffer was incubated in the chambers overnight at 4°C in a humidified chamber. Antibodies used were ID4 (1:200, Biocheck BCH9/82-12), α-SMA (1:100, Abcam ab5694), KRT14 (1:1000, Covance PRB-155P), KRT8 (1:500, DSHB TROMA1) and P63 (1:100, Novus NB100-691). The following day, slides were equilibrated to room temperature for 1-2 hr. Organoids were rinsed 2 times with IF buffer (0.1% BSA, 0.2% Triton-X-100, 0.05% Tween-20 in PBS) for 20 min each. Fluorescent secondary antibody (Jackson ImmunoResearch) diluted in blocking buffer (1:500) was added to the chambers and incubated for 45 min in the dark. Secondary antibody was rinsed off for 20 min in IF buffer, then 2 times with PBS for 10 min each. DAPI (Life Technologies) was incubated in the chambers for 15 min followed by a 10 min PBS rinse. The walls of the chamber slides were removed and one drop of Prolong Diamond mounting media (Thermo Fisher Scientific) was added to each well. Slides were coverslipped and edges sealed with clear nail varnish. Slides were allowed to dry for 24 hr at room temperature protected from light.

Organoids were imaged using a Leica DMI Sp8 confocal microscope using the 40X oil objective and 3X optical zoom. The FIJI software package was used to quantify the fluorescence within the organoids. For α-SMA, a circle was drawn around the entire organoid and fluorescence measured. For ID4, a mask was made using the DAPI channel and signal was measured within the nuclear mask. Multiple representative regions of background staining were quantified. Corrected fluorescence (CF) was calculated as described in (McCloy, Rogers et al., 2014). CF = integrated density – (area of selected organoid * mean fluorescence of all background readings)

### Cre mediated deletion of ID4 from primary cells

Following differential trypsinisation to remove irradiated 3T3 feeder cells, 1.0-1.5×10^5^ ID4floxGFP primary basal cells were seeded into a T75 with 7.5×10^5^ fresh irradiated 3T3 cells in 20 mL FAD media. Adenovirus (Vector Biolabs) was added to the media at a multiplicity of infection (MOI) of 100. Adenoviruses’ used were codon optimised Cre (iCre) and GFP adenovirus (#1772) and control eGFP adenovirus (#1060). Infection efficiency was assessed the following day using a fluorescence microscope to check GFP expression within the cells. Cells were harvested after 72 hr for downstream analysis.

### siRNA transfections

Comma-Dβ cells were seeded at a density of 1.5×10^4^ cells/cm^2^ into 6 well plates (Corning) for RNA-seq or 100 mm dishes (Corning) for ChIP-seq in antibiotic-free media. The following day siRNA constructs (Dharmacon) were transfected into the cells using Dharmafect-4 transfection reagent (Dharmacon) as per the manufacturer’s instructions at 20 nM. siRNA used were siGENOME Mouse Id4 SMARTpool (M-043687) and ON-TARGETplus Non-targeting Control Pool (D-001810). Media was changed the following day and cells were harvested 48 hr post-transfection.

### Western blotting

Cells were lysed in RIPA buffer (50 mM Tris-HCl pH 7.4, 1% NP-40, 0.5% Sodium Deoxycholate, 0.1% SDS, 1% Glycerol, 137.5 mM NaCl, 100 μM Sodium Orthovanadate, 20 μM MG132, 1 mM DTT and 1x cOmplete ULTRA Tablet (Roche)) and protein concentration was quantified using the Pierce BCA Protein Assay Kit (Thermo Fisher Scientific) according to the manufacturer’s instructions. Protein lysates (15-30 μg protein) were mixed with 1X NuPage loading buffer (Life Technologies) and 1X NuPage reducing agent (Life Technologies) and denatured by heating at 85°C for 5 min. Samples were run on a 4-12% Bis/Tris gels (Life Technologies) in MES or MOPS running buffer (Life Technologies). Protein was transferred to a 0.45 μm PVDF membrane (Merck Millipore) using BioRad transfer modules in transfer buffer (25 mM Tris, 192 mM Glycine, 20% Methanol). Membranes were blocked in Odyssey blocking buffer (LiCOR) for 1 hr at room temperature then incubated in primary antibody diluted in 5% BSA/TBS overnight at 4°C. The following antibodies were used for western blotting: ID4 (1:20,000, Biocheck BCH9/82-12), E2A (1:1000, from Dr Nicolas Huntington and Prof Lynn Corcoran from the Walter Eliza Hall Institute (WEHI)), HEB (1:1000, from Dr Nicolas Huntington and Prof Lynn Corcoran, WEHI or 1:1000 ProteinTech 144191-1-AP), Flag (1:5000, Sigma-Aldrich F1804), V5 (1:200, sc-58052), β-Actin (1:1000, Sigma-Aldrich A5441) and α-Tubulin (1:1000, Santa Cruz sc-5286). Fluorescent secondary antibody conjugated to IRDye680 or IRDye800 (LiCOR) diluted in Odyssey blocking buffer (1:15,000 - 1:20,000) were used for detection. An Odyssey CLx Infrared Imaging System (LiCOR) was used to image and quantify the western blots.

### Co-immunoprecipitation

Pierce Protein A/G magnetic breads (Thermo Fisher Scientific) were incubated with antibody for a minimum of 4 hr at 4°C on a rotating platform. For each IP 15 μL of beads and 2.5 μg antibody were used. Immunoprecipitating antibodies used were ID4 (pool of Santa Cruz sc-491 and sc-13047), E2A (from Dr Nicolas Huntington and Prof Lynn Corcoran, WEHI), HEB (Santa Cruz sc-357 and antibody from Dr Nicolas Huntington and Prof Lynn Corcoran, WEHI), Flag (Sigma-Aldrich F1804), V5 (Santa Cruz sc-58052 and Thermo Fisher R960-25), species matched IgG negative controls (sc-2027, sc-2025 and BioLegend 400602). Protein was extracted in IP lysis buffer (10% Glycerol, 0.03% MgCl_2_, 1.2% HEPES, 1% Sodium acid pyrophosphate, 1% Triton-X, 0.8% NaCl, 0.4% NaF, 0.04% EGTA, 1x complete ULTRA Tablet (Roche), 100 μM Sodium Orthovanadate, 20 μM MG132, 1 mM DTT and cells were passed 5 times through a 23 g needle to aid in lysis. Protein lysates (0.5-2 mg per IP) were added to washed beads and incubated overnight at 4°C on a rotating platform. Following washing, beads were resuspended in 2X NuPage loading buffer (Life Technologies) and 2X reducing agent (Life Technologies) in IP lysis buffer. Samples were incubated at 85°C for 5 min. Beads were separated on a magnetic rack and supernatant loaded onto NuPage gel for SDS-PAGE and western blotting. For detection of protein, fluorescent TrueBlot secondary antibodies were used (Jomar Life Research).

### RIME

The RIME protocol was adapted from the protocol developed by Mohammed *et al*. (Mohammed et al., 2013). Comma-Dβ cells were grown in 150 mm tissue culture dishes (Corning) until 80-90% confluent. A total of 8x 150 mm dishes were used per RIME sample. Cells were fixed with 1% PFA (ProSciTech) in DMEM/F12 (Gibco) for 7 min at room temperature on a rocking platform. Cross-linking was quenched by addition of molecular grade glycine (Astral Scientific) to a final concentration of 125 mM for 2 min on a rocking platform. Cells were washed twice with ice-cold PBS (Gibco) and scraped in 1 mL of ice-cold PBS containing magnesium and calcium salts (Gibco). Cells were centrifuged at 4°C for 5 min at 1200 rpm. Cell nuclei were enriched through a series of lysis buffers. Cells were first resuspended in 10 mL lysis buffer 1 (50 mM HEPES-KOH pH 7.5, 140 mM Sodium, 1 mM EDTA, 10% Glycerol, 0.5% NP-40 or Igepal CA-630, 0.25% Triton X-100, 100 μM Sodium Orthovanadate, 20 μM MG132, 1 mM DTT and 1x complete ULTRA Tablet (Roche) and incubated for 30 min at 4°C on a rotating platform. Cells were then centrifuged at 3000 g for 3 min at 4°C and supernatant removed. The pellet was then resuspended in 10 mL lysis buffer 2 (10 mM Tris-HCl pH 8.0, 200 mM NaCl, 1 mM EDTA, 0.5 mM EGTA, 100 μM Sodium Orthovanadate, 20 μM MG132, 1 mM DTT and 1x cOmplete ULTRA Tablet (Roche)) and incubated for a further 30 min at 4°C on a rotating platform. Following centrifugation, the resulting nuclei pellet was lysed in 2.5 mL lysis buffer 3 (10 mM Tris-HCl pH 8.0, 100 mM NaCl, 1 mM EDTA, 0.5 mM EGTA, 0.1% Sodium Deoxycholate, 0.5% N-lauroylsarcosine, 100 μM Sodium Orthovanadate, 20 μM MG132, 1 mM DTT and 1x complete ULTRA Tablet (Roche)). The nuclear lysate was sonicated using a Bioruptor sonicator (Diagenode) for 20 cycles of 30 sec on/30 sec off. Triton-X-100 was added to a final concentration of 1% and the sample was then centrifuged at 4700 g for 5 min at 4°C to remove cellular debris. Immunoprecipitation was carried out from the sheared nuclear supernatant described above with 20 μg of antibody and 100 μL of magnetic beads per sample. The following day, the beads were washed 10 times with RIPA buffer then 5 times with 100 mM ammonium hydrogen carbonate (AMBIC) (Sigma-Aldrich) solution to remove salts and detergents, resuspended in 50 μL AMBIC solution and transferred into clean tubes.

### Mass Spectrometry

Samples were processed as described in (Huang, Yang et al., 2015) for liquid chromatography coupled mass spectrometry (LC-MS/MS) using Sequential Windowed Acquisition of all THeoretical fragment-ion spectra (SWATH) acquisition. Briefly, samples were denatured in 100 mM triethylammonium bicarbonate and 1% sodium deoxycholate, disulfide bonds were reduced in 10 mM DTT, alkylated in 20 mM iodo acetamide, and proteins digested on the magnetic beads using trypsin. After C18 reversed phase (RP) StageTip sample clean up, peptides were analysed using a TripleToF 6600 mass spectrometer (SCIEX, MA, USA) coupled to a nanoLC Ultra 2D HPLC system (SCIEX). Peptides were separated for 60 min using a 15 cm chip column (ChromXP C18, 3 μm, 120 Å) (SCIEX) with an acetonitrile gradient from 3-35%. The MS was operated in positive ion mode using either a data dependent acquisition method (DDA) or SWATH acquisition mode. DDA was performed of the top 20 most intense precursors with charge stages from 2+ - 4+ with a dynamic exclusion of 30 s. SWATH-MS was acquired using 100 variable size precursor windows. DDA files were searched using ProteinPilot 5.0 (SCIEX) against the reviewed UniProt *Mus musculus* protein database (release February 2016) using an unused score of 0.05 with decoy search strategy enabled. These search outputs were used to generate a spectral library for targeted information extraction from SWATH-MS data files using PeakView v2.1 with SWATH MicroApp v2.0 (SCIEX) importing only peptides with < 1% FDR. Protein areas, summed chromatographic area under the curve of peptides with extraction FDR ≤ 1%, were calculated and used to compare protein abundances between bait and control IPs.

### ChIP-seq

Our ChIP-seq protocol was adapted from (Khoury, Achinger-Kawecka et al., 2020). Comma-Dβ cells were grown in 150 mm or 100 mm dishes (Corning) until 80-90% confluent. For the ChIP-seq experiment on unperturbed cells 2x 150 mm dishes were used for HEB ChIP and 1x 150 mm dishes were used for histone marks. For the ID4 siRNA experiment 3x 100 mm dishes per condition were used. Cells were scraped in ice-cold PBS (Gibco) and resuspended by passing 10 times through a 19 g syringe. Cells were fixed in 1% PFA (ProSciTech) in PBS (Gibco) for 15 min at room temperature followed by quenching with glycine to a final concentration of 125 mM for 5 minutes followed by two PBS washes. Nuclei was extracted by resuspending cells in nuclei extraction buffer (10 mM Tris-HCl pH 7.5, 10 mM NaCl, 3 mM MgCl_2_, 0.1 mM EDTA, 0.5% IGEPAL, 1x cOmplete ULTRA Tablet (Roche)) and incubated on ice for 10 min followed by 20 passes on a tight Dounce homogeniser, or until nuclei were extracted. Nuclei were visually inspected using trypan blue and a haemocytometer. Nuclei were pelleted by centrifugation and resuspended in sonication buffer (50 mM Tris-HCl pH 8, 1% SDS, 10 mM EDTA, 1x cOmplete ULTRA Tablet (Roche)) and sonicated using a Bioruptor sonicator (Diagenode) to achieve DNA fragments between 100-500 bp with a mean fragment size of approximately 300 bp. In the siRNA experiment, protein was quantified using the Pierce BCA assay to load equal amounts of input material for NT and ID4 siRNA for immunoprecipitation. Sonicated sample was diluted with IP dilution buffer to 1 mL (16.7 mM Tris-HCl pH 8, 0.01% SDS, 1% Triton X-100, 167 mM NaCl, 1.2 mM EDTA). Samples were cleared with protein A/G magnetic beads for 1.5 hr at 4°C. 1% of the cleared nuclear lysate was removed for the input control and immunoprecipitation was performed as described previously. For HEB ChIP 100 μL of beads and 20 μg of HEB antibody (sc-58052) was used with chromatin from approximately 20-30×10^6^ cells. For histone mark ChIP, 50 μL beads with 10 μg of the following antibodies – H3K4Me3 (Active Motif 28431), H3K27Me3 (Merck Millipore 07-449), was used with chromatin from approximately 10-15×10^6^ cells. The following day, beads were washed for 5 min each with 1 mL of the following buffers: Low salt buffer (0.1% SDS, 1% Triton X-100, 2 mM EDTA, 20 mM Tris-HCl, pH 8, 150 mM NaCl), High salt buffer (0.1% SDS, 1% Triton X-100, 2 mM EDTA, 20 mM Tris-HCl, pH 8, 500 mM NaCl), LiCl buffer (0.25 M LiCl, 1% IGEPAL, 1% deoxycholic acid (sodium salt), 1 mM EDTA, 10 mM Tris, pH 8) followed by 2 washes with TE buffer (10 mM Tris-HCl, 1 mM EDTA, pH 8). DNA was eluted twice in 100 uL ChIP elution buffer (1% SDS, 0.1 M Sodium Biocarbonate) for 15 minutes at room temperature each. Cross-linking was reversed overnight by treating samples with 200 mM NaCl and 250 μg/mL Proteinase K (New England Biolabs) and incubating overnight at 65°C. The following day samples were treated with 100 μg/mL RNAse A (Qiagen) for 1 hr at 37°C. DNA was purified using Phase Lock Gel Light tubes (Quantabio 5Prime) according to the manufacturer’s instructions. DNA was eluted in 20 uL nuclease-free TE buffer, pH8 (Qiagen).

DNA concentration was measured using the Qubit HS Assay Kit (Thermo Fisher Scientific). Libraries were prepared using the Illumina TruSeq ChIP library prep kit (Illumina) following the manufacturer’s instructions except the gel purification step was replaced with a two-sided AMPure XP bead (Beckman Coulter) size selection to obtain libraries between 200-500 bp. Library sizes were verified using the 4200 TapeStation System (Agilent) with a D1000 ScreenTape (Agilent) and concentration was determined using the Qubit HS Assay Kit (Thermo Fisher Scientific). ChIP libraries were sequenced on the NextSeq system (Illumina), with 75 bp paired-end reads.

Reads were aligned with BWA (Li & Durbin, 2009) and all reads with a MAPQ<15 were removed. Alignment statistics were generated using Samtools Flagstat (Li, Handsaker et al., 2009). ChIP-seq peaks were called using the peak calling algorithm MACS (Zhang, Liu et al., 2008) and ENCODE blacklist regions were removed. For unperturbed cells, consensus peaks present in at least 2 of 4 replicates were used for downstream analysis. Due to lower amount of input material available from the siRNA experiment, less consensus peaks were called and for this reason analysis was conducted on merged peaks from the 3 replicates. Motif enrichment analysis was performed using MEME-ChIP (Machanick & Bailey, 2011). Peaks were annotated to genomic features using HOMER (Heinz, Benner et al., 2010). GREAT was used for functional enrichment analysis (v4.0.4) and gene annotation using the default parameters (McLean et al., 2010). SeqPlots (Stempor & Ahringer, 2016) and IGV (Robinson, Thorvaldsdóttir et al., 2011) software were used for data visualisation. Differential binding analysis was performed using the DiffBind package (Stark, 2011). For overlapping of of siID4 RNA-seq and HEB ChIP-seq genes, a hypergeometric test was used to determine if overlap was significant, assuming 30,000 genes in the mouse genome.

## Data availability

The mass spectrometry proteomics data have been deposited to the ProteomeXchange Consortium via the PRIDE partner repository with the dataset identifier PXD017517.

Processed data files for the RNA-seq (counts tables) and ChIP-seq (bed files) have been uploaded as Figure Source Data. Raw data files are in the process of being uploaded onto the GEO public repository and will be made available prior to publication. We can provide the raw data files upon request.

## Acknowledgements

We are very grateful for Amanda Khoury for her extensive advice on the ChIP-seq experiments, Samantha Oakes for providing the mammary gland FFPE blocks from different developmental stages, David Gallego Ortega for advice on tissue dissociation and sorting and William Hughes for help with microscopy image analysis. We thankfully acknowledge the support of the Australia Postgraduate Award and The National Health and Medical Research Council of Australia (1107671). Aspects of this research were supported by access to the Australian Proteome Analysis Facility, funded by the Australian Government’s National Collaborative Research Infrastructure Scheme. T.R.C and J.N.S are supported by the Cancer Institute NSW (171105) and a Career Catalyst grant from Susan G. Komen.

## Author Contributions

H.H designed and performed the experiments and wrote the manuscript. D.R analysed the RNA-seq and ChIP-seq data. S.J co-supervised the project, contributed to experiments and discussions and extensively revised the manuscript. S.Z.W analysed public scRNA-seq data and revised the manuscript. L.A.B advised on RIME experiments and contributed to discussions. C.K and M.P.M performed mass spectrometry and analysis. C.H performed RNA-seq library preps and analysis. A.M assisted with mouse experiments. J.N.S and T.R.C performed picrosirius red image analysis. B.P assisted with ChIP-seq experiments. N.H contributed IP antibodies. C.J.O co-supervised the project and revised the manuscript. J.S.C helped to conceptualise the project and revised the manuscript. J.V contributed the reporter mice and revised the manuscript. A.S conceived the study and was in charge of overall direction and planning.

## Supplementary Figure legends

**Figure S1 related to Figure 1.**

**a)** Proportions of basal, luminal progenitor (LP), and mature luminal (ML) subpopulations between ID4 WT and KO mammary epithelial cells. Unpaired two-tailed students t-test. ns = not significant. Error bars represent SEM. N=4. Representative GSEA enrichment plots displaying the profile of the running Enrichment Score (green) and positions of gene set members on the rank ordered list for pathways related to growth **(b)** and myoepithelial function **(c).**

**Figure S2 related to Figure 2.**

**a)** Co-immunofluorescent staining of ID4 and α-SMA in a section containing both a TEB and a duct from a pubertal (6 week) mammary gland. Scale bar = 20 μm. **b)** Mouse mammary glands at different developmental stages were stained by IHC for ID4. Scale bar = 100 μm. Representative images from 3 animals per stage shown. High power insets are displayed below images. **c)** Differentiation trajectory of basal cells from (Bach et al., 2017) coloured by developmental stage (upper) and pseuodotime (lower). Low values (dark blue) represent undifferentiated cells. NP = Nulliparous (8 week), G= Gestation (Day 14.5), L = Lactation (Day 6), I = Involution (Day 11). **d)** Expression of *Id4, Cnn1, Myh11, Acta2, Mylk* and *Oxtr* over pseudotime.

**Figure S3 related to Figure 3.**

**a)** Phase contrast images of passage 11 conditionally reprogrammed cells grown in the presence and absence of ROCK inhibitor Y-27632, irradiated NIH-3T3 feeder cells, and Matrigel (MG) coated tissue culture flasks. Scale bar = 100 μm. b) ID4 western blot in cell lysates collected from cells shown in panel (a). ID4 from basal cells runs at a higher molecular weight to ID4 expressed by NIH-3T3 cells. **c)** Co-immunofluorescent staining of primary basal cells with basal markers P63 and KRT14. Scale bar = 20 μm. **d)** Basal cells grown in 3D as organoids on top a plug of Matrigel. Confocal image of an organoid stained for luminal marker KRT8 and basal marker KRT14. Scale bar = 10 μm.

**Figure S4 related to Figure 4.**

**a)** Genes were ranked based on the limma t-statistic comparing NT and siID4 cells and GSEA was carried out using the Hallmark gene sets. The top 10 positively (red) and negatively (blue) enriched pathways are displayed. **b)** Representative Hallmark GSEA enrichment plots displaying the profile of the running Enrichment Score (green) and positions of gene set members on the rank ordered list.

**Figure S5 related to Figure 5.**

Co-immunoprecipitation and western blotting of ID4, E2A and HEB from human normal-like mammary epithelial cell lines PMC42 (a) and MCF10A cells overexpressing Flag-tagged ID4 protein **(b). c)** Unsupervised hierarchical clustering heat map of SWATH RIME data from Comma-Dβ cells. Proteins with significantly higher abundance (p-value<0.05) in the HEB IPs compared to IgG IPs are displayed. Log2 protein area was used to generate the heatmap.

**Figure S6 related to Figure 6.**

**a)** Genomic distribution of HEB consensus peaks. TSS = transcriptional start site. TTS = transcriptional termination site. **b)** Heatmaps of HEB, H3K4Me3, H3K27Ac, and H3K27Me3 ChIP-seq signal at HEB-bound regions. **c)** Examples of HEB and histone mark peaks occurring upstream of ID4 repressed genes *Col1a2, Col3a1*, and *Col5a1* (multiple HEB peaks) from the Integrative Genomics Viewer (IGV). Bars beneath peaks represent consensus MACS call (FDR<0.05) in at least 2 of 4 biological replicates. Input was used as a negative control. Purple boxes highlight HEB binding regions. Refseq genes shown in blue. Data scales for each track are indicated. **d)** Western blot analysis of ID4 and HEB expression in Comma-Dβ cells treated with NT or ID4-targeting siRNA. Densitometry quantification of ID4 and HEB bands. Band intensity was normalised to β-Actin and expressed as fold change relative to NT control. N=3. Unpaired two-tailed students t-test. **** p<0.0001. ns = not significant Error bars represent SEM. **e)** Top 4 enriched transcription factor binding motifs determined using MEME-ChIP for consensus HEB ChIP-seq peaks in Comma-Dβ cells treated with NT or ID4-targeting siRNA. E-values are displayed. **f)** Example of HEB peaks in control NT cells and siID4 cells upstream of *Col5a1* from the Integrative Genomics Viewer (IGV). Green boxes highlight HEB binding regions. Individual replicates and merged tracks are displayed. Refseq genes shown in blue. Data scales indicated, g) GREAT pathway analysis of 263 peaks increased in siID4 compared to NT from Fig. 6f. Top 16 Gene Ontologies (Biological process, cellular component, molecular function) are displayed.

## Supplementary Tables

**Table S1.** All differentially expressed genes between ID4 wild type and knockout sorted mammary populations (Basal, luminal progenitor and mature luminal).

**Table S2.** Top 50 differentially expressed genes between *Id4*-high and *Id4-lo* basal cells from Bach et al. single cell RNA-seq analysis.

**Table S3.** Top 50 differentially expressed genes between siID4 and Non-targeting control Comma-Dβ cells.

**Table S4.** Proteins identified in ID4 and HEB RIME experiments.

**Table S5.** Genomic regions of Differentially bound HEB ChIP-Seq peaks between siID4 and NT control Comma-Dβ cells.

