## Supplementary figures and images for "Inhibitor of Differentiation 4 (ID4) represses myoepithelial differentiation of mammary stem cells through its interaction with HEB"

### Figure S1

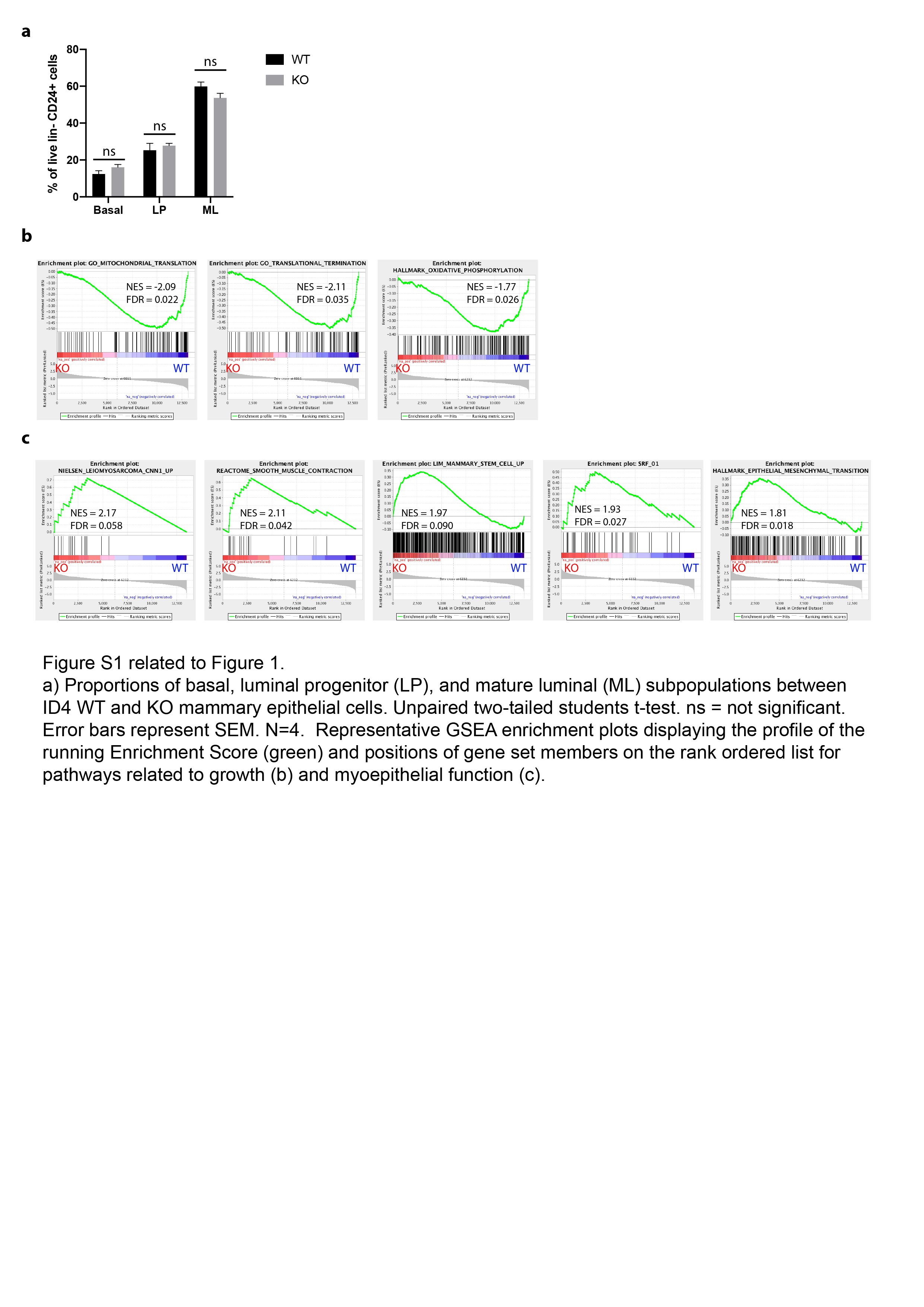

### Figure S2

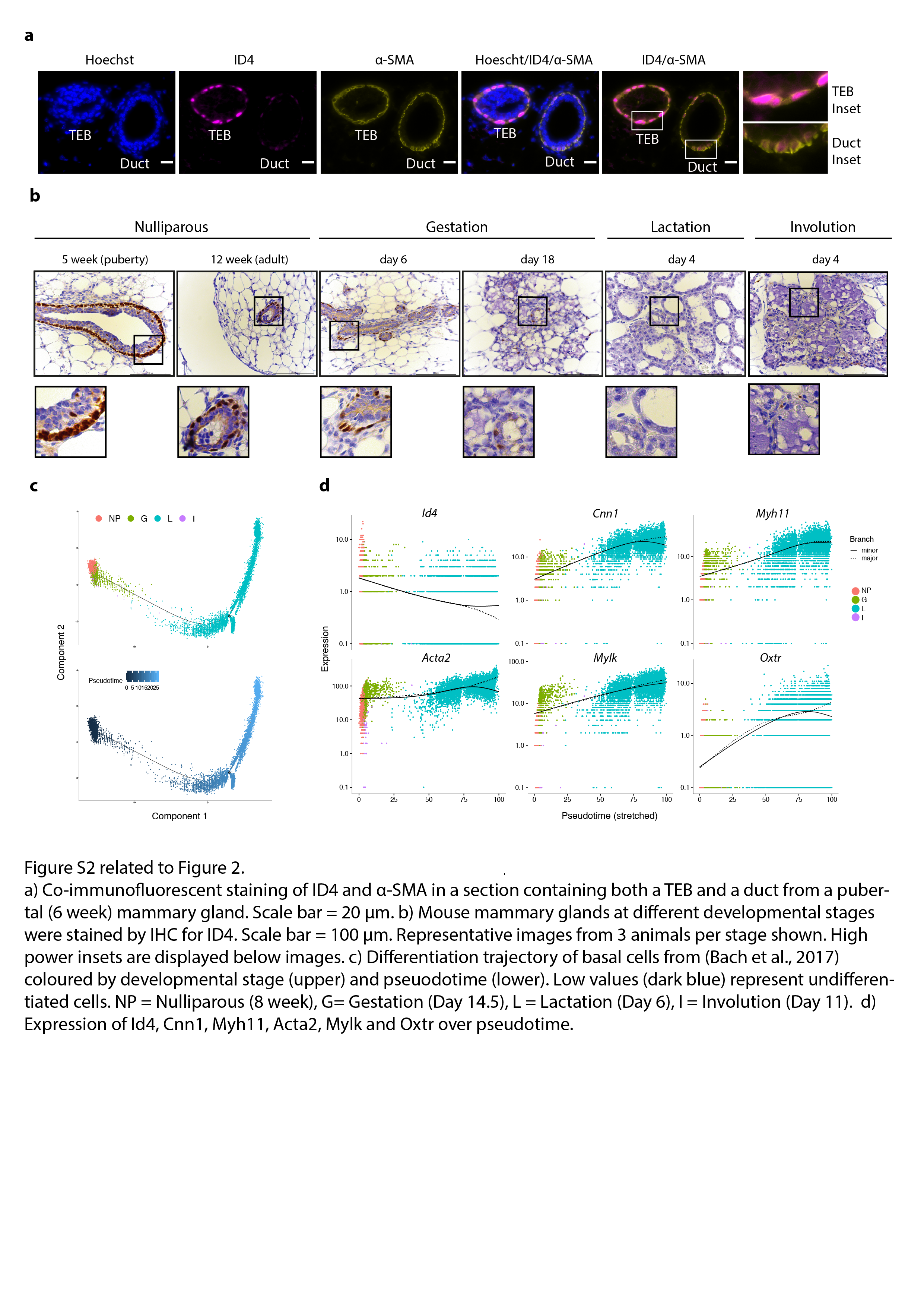

### Figure S3

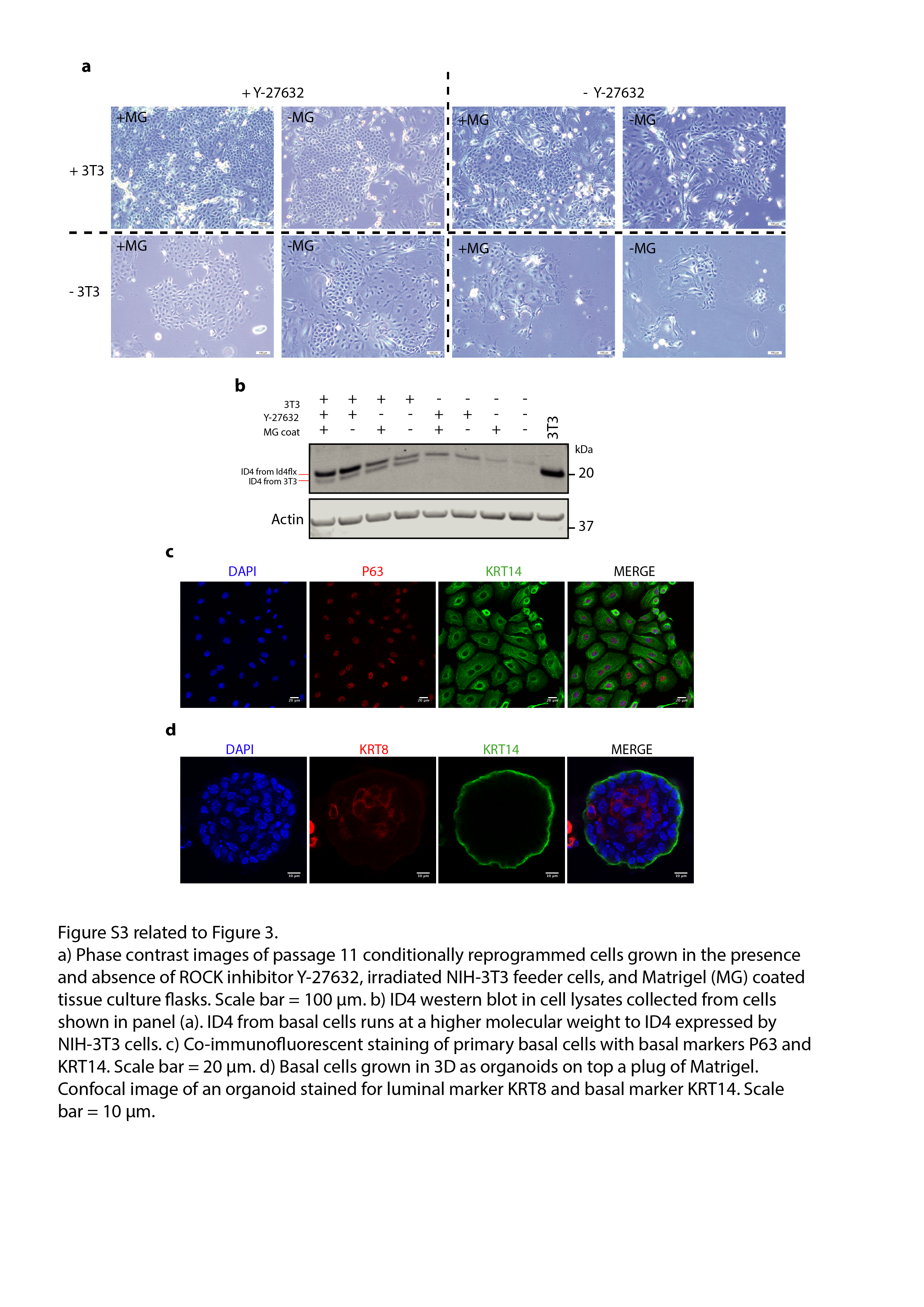

### Figure S4

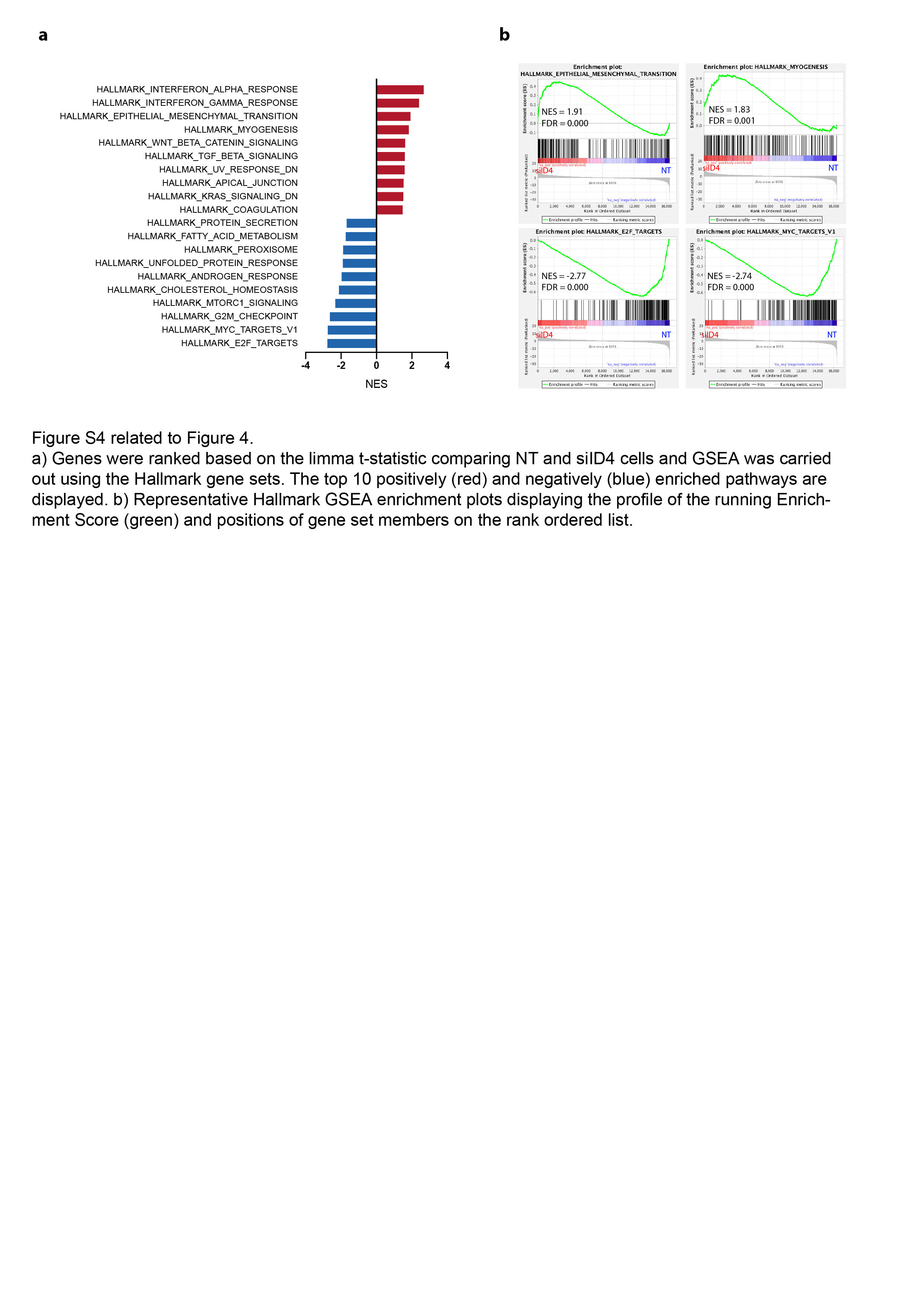

### Figure S5

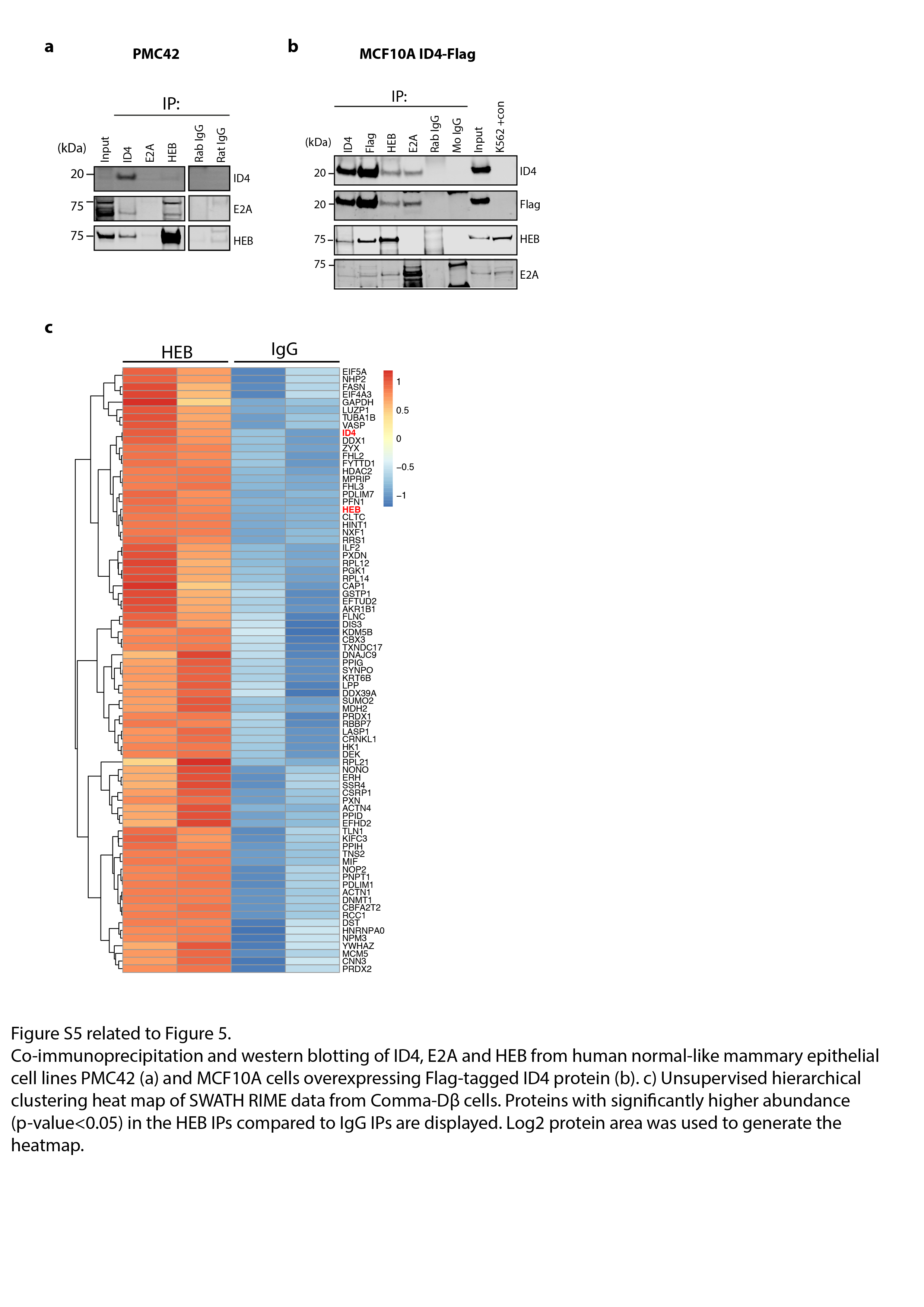

### Figure S6

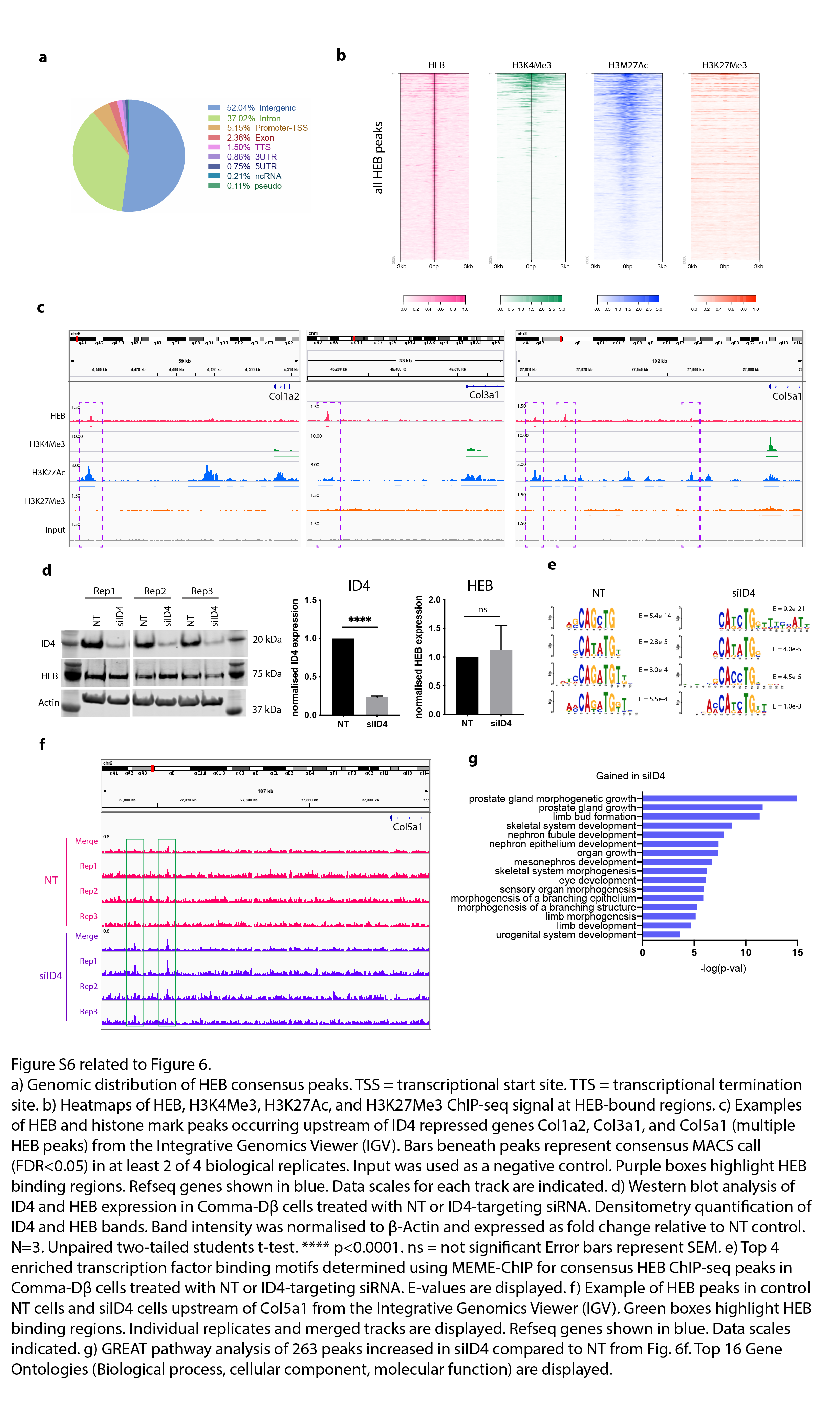
