## Supplementary material for "Inhibitor of Differentiation 4 (ID4) represses myoepithelial differentiation of mammary stem cells through its interaction with HEB": Tables S1-5

**Table S1.** Differentially expressed genes between ID4 wild type and knockout sorted mammary populations (Basal, luminal progenitor and mature luminal).

Down in ID4 KO Basal cells

| Symbol | logFC | FDR |
|---|---|---|
| Slain1 | -5.8761593 | 1.16E-07 |
| Tec | -5.4685621 | 6.90E-06 |
| Rab25 | -3.9419651 | 4.82E-05 |
| Gm37249 | -7.6807283 | 9.06E-05 |
| Slc37a1 | -5.7836186 | 0.00025583 |
| Mrps26 | -4.902518 | 0.00033316 |
| Spint1 | -2.1929813 | 0.00033316 |
| Vipr1 | -4.1040324 | 0.00075387 |
| Rnase1 | -8.4173066 | 0.00102459 |
| Gm38380 | -8.0127451 | 0.00159031 |
| Rasef | -2.3528078 | 0.00159031 |
| Whamm | -3.3625247 | 0.00172406 |
| Pfkfb2 | -2.0966066 | 0.00172406 |
| Ccdc149 | -6.3017489 | 0.00180465 |
| Esrrg | -4.1412709 | 0.00245307 |
| Pitpnm3 | -3.9308061 | 0.00450314 |
| Pik3cg | -4.421759 | 0.00450425 |
| Zdhhc14 | -3.2550828 | 0.00450425 |
| 2310001H17Rik | -4.634897 | 0.00502286 |
| Ero1lb | -2.4799475 | 0.00502286 |
| 1700019G17Rik | -5.1958717 | 0.00524307 |
| Cadps2 | -2.2214832 | 0.00524307 |
| Lcp2 | -10.809021 | 0.0053489 |
| Bdh1 | -4.7242882 | 0.0053489 |
| Pde7b | -2.4765853 | 0.0053489 |
| Pgap2 | -3.2132228 | 0.0058085 |
| Dffa | -3.0124642 | 0.0058085 |
| Gm17638 | -3.8568908 | 0.00607114 |
| Cdo1 | -2.2295634 | 0.00696051 |
| Gm9930 | -3.6358181 | 0.00735289 |
| Gm26816 | -4.8519903 | 0.00789403 |
| Elovl7 | -3.0621941 | 0.00856546 |
| Gm36970 | -7.4055617 | 0.00889776 |
| Cd160 | -3.6195487 | 0.00889776 |
| Fadd | -2.5753838 | 0.00889776 |
| Rbck1 | -1.9398277 | 0.00899399 |
| Gm13590 | -6.7229983 | 0.00956316 |
| Sema4d | -2.2331352 | 0.00966214 |
| Cyp2f2 | -6.0620549 | 0.01055025 |
| Hmgcs2 | -3.0871286 | 0.01055025 |
| 4930448N21Rik | -6.0646566 | 0.01056653 |
| Dennd6b | -3.9906293 | 0.01056653 |
| Esr1 | -1.8071592 | 0.01056653 |
| Nedd4l | -1.480922 | 0.01088434 |
| RP23-441I24.5 | -7.2551155 | 0.01107342 |
| F5 | -7.4192872 | 0.0116144 |
| Prr18 | -6.7460573 | 0.0116144 |

Up in ID4 KO Basal cells

| Symbol | logFC | FDR |
|---|---|---|
| H2-Q6 | 4.31613774 | 4.75E-05 |
| Id3 | 2.63682912 | 9.06E-05 |
| Ano10 | 3.89804479 | 0.00046174 |
| Tmem170 | 2.30862261 | 0.00068512 |
| Ogfod2 | 4.22077896 | 0.00068512 |
| Gm37699 | 5.43072795 | 0.00081392 |
| Tnfaip6 | 7.80941583 | 0.00142474 |
| Pcdhga2 | 9.19841087 | 0.00142474 |
| Zbed3 | 2.26298816 | 0.00327707 |
| Pih1d1 | 4.21931897 | 0.00450425 |
| Bcan | 7.63505091 | 0.00450425 |
| Ell3 | 4.24504233 | 0.00502286 |
| Zfp563 | 6.38608928 | 0.00708898 |
| Rps6ka1 | 1.71722808 | 0.00765046 |
| Rpusd2 | 3.61285965 | 0.00765046 |
| Fhod1 | 3.94570455 | 0.00765046 |
| Zc3h6 | 2.9195023 | 0.00797368 |
| Dhrs7 | 3.26418384 | 0.00816478 |
| Scd3 | 7.19641963 | 0.00856546 |
| Gm11651 | 8.5030334 | 0.00856546 |
| 2810408I11R | 6.24222871 | 0.00866565 |
| Zeb2 | 3.26379968 | 0.00889776 |
| Fancb | 7.49065765 | 0.00956316 |
| Tcf4 | 1.13399286 | 0.00966214 |
| Itih2 | 3.40864043 | 0.00966214 |
| Slc14a1 | 3.67730636 | 0.01055025 |
| F8a | 6.500476 | 0.0105759 |
| Gm29358 | 3.07990109 | 0.01236038 |
| Lrrc56 | 7.12013439 | 0.0138148 |
| Zfp775 | 7.37409308 | 0.01393153 |
| Slc12a6 | 1.93113856 | 0.01647764 |
| Gm14005 | 2.77591451 | 0.01731133 |
| Slc7a4 | 3.93057143 | 0.01731133 |
| Nup133 | 2.76476939 | 0.01846882 |
| Mus81 | 3.66199138 | 0.0189877 |
| Smarcd2 | 2.4679476 | 0.01965463 |
| Pqlc2 | 6.78969776 | 0.0205907 |
| Gm14133 | 7.28323154 | 0.02269248 |
| 2610305D13 | 3.8400096 | 0.02459026 |
| Gm20632 | 3.24784473 | 0.0246672 |
| Cox17 | 2.37316844 | 0.02551131 |
| Figf | 6.24705504 | 0.0260315 |
| Kcnj15 | 3.97985264 | 0.02682117 |
| Chchd3 | 1.69661482 | 0.0274508 |
| Ankfy1 | 1.20166358 | 0.02938045 |
| Sepsecs | 2.32268549 | 0.02938045 |
| Lgals9 | 1.42214532 | 0.02946512 |

| | | |
|---|---|---|
| RP23-451E8.7 | -5.8737876 | 0.01202769 |
| Lsm10 | -5.8209182 | 0.01202769 |
| Gm37678 | -6.8182812 | 0.01234847 |
| Nos1ap | -2.1288097 | 0.01236038 |
| Psme3 | -1.2383358 | 0.01236038 |
| Gab2 | -3.0827818 | 0.0127676 |
| RP23-331L5.1 | -8.4069986 | 0.01335499 |
| Zfp763 | -8.2836233 | 0.01393153 |
| 2410015M20Rik | -2.9896988 | 0.01447117 |
| Sdr42e1 | -3.0082093 | 0.01469589 |
| Sh3yl1 | -3.8049769 | 0.01506439 |
| Gm26668 | -5.8673695 | 0.01605985 |
| Zfp750 | -3.200232 | 0.01653997 |
| Gm5124 | -6.0586427 | 0.01654103 |
| Nuak2 | -3.4166389 | 0.01791017 |
| Neu1 | -2.2798247 | 0.01791017 |
| Ergic2 | -1.299788 | 0.01846882 |
| Irf4 | -5.4685664 | 0.01909823 |
| Susd4 | -6.4930829 | 0.0191241 |
| Eif3b | -1.1949702 | 0.0191241 |
| Zswim5 | -5.3455119 | 0.01965463 |
| Zfp747 | -2.5843511 | 0.01965463 |
| Emc4 | -2.3283571 | 0.01988716 |
| S100a8 | -6.0326537 | 0.02018108 |
| Tmem218 | -4.0102175 | 0.02018108 |
| RP24-555P13.4 | -8.5139411 | 0.02061243 |
| Lrrc48 | -6.9678476 | 0.02061243 |
| Gm17137 | -2.8841532 | 0.02269248 |
| Gm26511 | -7.0979157 | 0.02308312 |
| Tubd1 | -6.9298202 | 0.02464107 |
| Rhov | -2.4906502 | 0.02551131 |
| Serpinb1b | -9.5864184 | 0.02578832 |
| A430005L14Rik | -6.8314304 | 0.02578832 |
| Gm28036 | -9.0796576 | 0.02581414 |
| Kbtbd8 | -6.255735 | 0.02581414 |
| Gm29670 | -6.2099579 | 0.02585538 |
| Kcnj9 | -7.622608 | 0.02617521 |
| Tmem8 | -5.7353159 | 0.02682117 |
| Tmem144 | -4.7865513 | 0.02682117 |
| Slmo2 | -1.7032178 | 0.02682117 |
| C430002E04Rik | -7.2647216 | 0.0270328 |
| Dnase1l1 | -3.3423945 | 0.02892618 |
| Rrp12 | -2.2959923 | 0.02892618 |
| Tnfaip2 | -2.9333162 | 0.02938045 |
| Cd3e | -7.4187097 | 0.02946512 |
| Dlg5 | -1.2377106 | 0.02946512 |
| Cenpb | -1.809591 | 0.03145902 |
| Zfp595 | -3.6085529 | 0.03179418 |
| B130006D01Rik | -4.3509367 | 0.03206114 |
| Arrdc1 | -2.053794 | 0.03206114 |
| E130208F15Rik | -6.6531997 | 0.03231049 |
| Gm14227 | -5.7790586 | 0.03235502 |

| | | |
|---|---|---|
| Mfi2 | 3.73649745 | 0.02946512 |
| Fut2 | 6.94446871 | 0.02946512 |
| Atg9a | 1.90820942 | 0.03022162 |
| Wdr90 | 2.97089881 | 0.03034741 |
| RP23-7E4.4 | 5.81007305 | 0.03034741 |
| Zfp874b | 4.15882277 | 0.03050578 |
| Zfp780b | 3.19152822 | 0.03179418 |
| Ebf3 | 4.97524464 | 0.03179796 |
| Ly6a | 1.76508176 | 0.03205577 |
| Ext2 | 2.03460945 | 0.03205577 |
| Mpdu1 | 2.86206202 | 0.03205577 |
| March9 | 3.83478776 | 0.03205577 |
| Slc35d3 | 7.81702611 | 0.03205577 |
| Ajap1 | 8.08721944 | 0.03235502 |
| 4930515G01 | 8.56109828 | 0.03263143 |
| Tmem54 | 4.76727195 | 0.03270391 |
| Gm38042 | 6.07150199 | 0.03270391 |
| Hcar2 | 3.34600908 | 0.03398368 |
| Sytl1 | 3.61906416 | 0.03417395 |
| RP24-188G23 | 5.93916928 | 0.03417395 |
| Paox | 5.02771966 | 0.03505378 |
| Zeb1 | 2.99668138 | 0.03569185 |
| Gm26587 | 8.97300958 | 0.03656874 |
| RP23-218L17 | 6.74341532 | 0.03691029 |
| Gm6498 | 7.65016491 | 0.03698683 |
| Ebf2 | 5.95366037 | 0.03717174 |
| Gimap4 | 7.82755036 | 0.03739897 |
| Slfn10-ps | 3.10650374 | 0.0376935 |
| Trim34a | 4.57716303 | 0.0390837 |
| Csgalnact1 | 3.92564714 | 0.03910804 |
| Maoa | 1.22511263 | 0.03931921 |
| Hist1h1d | 6.96372886 | 0.04176468 |
| Aldh3b1 | 4.77564281 | 0.04200334 |
| Tubg2 | 4.75380928 | 0.04325084 |
| Ambra1 | 1.62988888 | 0.04329228 |
| Erich1 | 4.46923584 | 0.04329228 |
| Klhl12 | 1.67478222 | 0.04520409 |
| Tmem256 | 3.44093779 | 0.04520409 |
| Slc22a18 | 5.32676839 | 0.04520409 |
| Gm13212 | 6.51259179 | 0.04604043 |
| 0610009B22I | 2.16752529 | 0.04635119 |
| Adgrd1 | 4.70771842 | 0.04760645 |
| Gm15539 | 7.14761049 | 0.04845168 |
| Pi16 | 4.54199869 | 0.04888851 |
| Ppm1l | 1.26829856 | 0.04906974 |
| Gosr1 | 2.44997811 | 0.04906974 |
| Ephx2 | 5.44973843 | 0.04906974 |
| Rbp4 | 5.83152068 | 0.04925528 |
| Igfbp6 | 6.33730028 | 0.04925955 |

| | | |
|---|---|---|
| Gm12403 | -6.0820602 | 0.03639202 |
| Kcnab3 | -7.4735576 | 0.03656874 |
| Mthfsl | -2.9279644 | 0.03698683 |
| Dclre1b | -2.6239393 | 0.03698683 |
| Manba | -2.2533337 | 0.03698683 |
| Gm37945 | -7.5598995 | 0.0376935 |
| Sbsn | -3.5801147 | 0.0376935 |
| Anxa4 | -1.0912073 | 0.0376935 |
| F2rl2 | -4.3242468 | 0.03792079 |
| Gm16144 | -3.4590156 | 0.03792079 |
| Trim7 | -2.3908693 | 0.03910804 |
| Tmem241 | -3.0634371 | 0.04038653 |
| Gpx3 | -2.0126216 | 0.0409149 |
| Fam174b | -2.555321 | 0.04231119 |
| Extl1 | -4.1801469 | 0.04248817 |
| Top3a | -2.5768171 | 0.04248817 |
| Arhgef37 | -6.4817292 | 0.04329228 |
| Chil1 | -1.4544817 | 0.04329228 |
| Rnf169 | -1.1577514 | 0.04329228 |
| Ift43 | -1.8368361 | 0.04441894 |
| | -1.5795483 | 0.04482725 |
| Olr1 | -8.4182487 | 0.04520409 |
| Asic1 | -5.5331749 | 0.04520409 |
| 1700007E05Rik | -3.7986006 | 0.04520409 |
| Necap1 | -2.3099298 | 0.04520409 |
| Isg20 | -2.1765914 | 0.04520409 |
| Plekhm1 | -1.2081944 | 0.04520409 |
| Gm9874 | -3.288193 | 0.04629926 |
| Gm26593 | -5.3423678 | 0.04634547 |
| Sdhaf1 | -4.3381042 | 0.04635119 |
| Eci1 | -2.799025 | 0.0463883 |
| Mapre3 | -2.6726199 | 0.0463883 |
| Smim24 | -4.7774515 | 0.04733769 |
| Gstt2 | -4.309634 | 0.04784973 |
| Slc22a4 | -5.5639502 | 0.04845168 |
| Gm37289 | -6.7723859 | 0.04906974 |
| 9330159N05Rik | -5.9951296 | 0.04906974 |
| Rab17 | -4.3010224 | 0.04906974 |
| Iba57 | -4.2425044 | 0.04906974 |
| Mrpl14 | -3.1973962 | 0.04906974 |
| D6Wsu163e | -2.4092623 | 0.04906974 |
| Trmt10a | -1.9001821 | 0.04906974 |
| Mgst3 | -2.8681105 | 0.04933717 |

LP DEGs

| Symbol | log FC | FDR |
|---|---|---|
| Dst | -10.722656 | 0.00119204 |
| Dst | 9.02219776 | 0.00332346 |

ML DEGs

| Symbol | logFC | FDR |
|---|---|---|
| Pappa | -3.5149615 | 2.50E-07 |
| Pdk4 | -2.0713096 | 0.03540697 |

**Table S2.** Top 50 differentially expressed genes between *Id4* -high and *Id4* -lo basal cells from Bach et al. single cell RNA-seq analysis.

| Gene | logFC | p_val_adj | cluster |
|---|---|---|---|
| Id4 | 2.103 | 2.7E-278 | High |
| Slpi | 1.967 | 1.1E-09 | High |
| Ncl | 1.053 | 3.9E-47 | High |
| Mthfd2 | 0.983 | 7.6E-27 | High |
| Igfbp3 | 0.957 | 9.3E-15 | High |
| Ctsl | 0.938 | 7.0E-35 | High |
| Nme1 | 0.930 | 4.5E-34 | High |
| Nars | 0.907 | 1.8E-41 | High |
| Gnl3 | 0.890 | 1.2E-27 | High |
| Aqp3 | 0.858 | 2.8E-03 | High |
| Fst | 0.858 | 1.4E-13 | High |
| Eif4ebp1 | 0.835 | 2.6E-34 | High |
| Srsf2 | 0.832 | 1.9E-38 | High |
| Rsl1d1 | 0.811 | 4.8E-34 | High |
| Eif4a1 | 0.783 | 3.4E-40 | High |
| Rrs1 | 0.778 | 7.6E-24 | High |
| Nupr1 | 0.775 | 2.0E-16 | High |
| Ddx21 | 0.772 | 2.6E-29 | High |
| Arl4c | 0.766 | 2.6E-25 | High |
| Gars | 0.766 | 1.5E-19 | High |
| Eif2s2 | 0.762 | 3.1E-37 | High |
| Ran | 0.758 | 1.0E-41 | High |
| Nhp2 | 0.747 | 4.3E-24 | High |
| Nop56 | 0.741 | 6.5E-25 | High |
| Anp32b | 0.735 | 3.3E-38 | High |
| Cxcl1 | 0.733 | 6.8E-10 | High |
| Sfn | 0.729 | 2.7E-19 | High |
| Srm | 0.725 | 2.0E-25 | High |
| Gjb4 | 0.720 | 5.2E-12 | High |
| Eif5a | 0.714 | 4.4E-33 | High |
| Epcam | 0.714 | 1.3E-18 | High |
| Snrpd1 | 0.709 | 3.6E-25 | High |
| Tars | 0.708 | 2.3E-22 | High |
| Sdc1 | 0.703 | 2.7E-19 | High |
| Herpud1 | 0.688 | 1.3E-25 | High |
| Cotl1 | 0.685 | 3.7E-35 | High |
| Tnfrsf12a | 0.682 | 1.8E-24 | High |
| Plaur | 0.674 | 1.4E-17 | High |
| Srsf3 | 0.672 | 4.2E-34 | High |
| Cda | 0.669 | 2.4E-17 | High |
| Lgals3 | 0.662 | 1.5E-09 | High |
| Hspa9 | 0.656 | 3.3E-29 | High |
| Eif3g | 0.656 | 3.1E-28 | High |
| Nolc1 | 0.651 | 3.1E-23 | High |

| Gene | logFC | p_val_adj | cluster |
|---|---|---|---|
| Fos | 1.482 | 1.4E-39 | Low |
| Egr1 | 1.427 | 5.7E-35 | Low |
| Jund | 1.388 | 3.3E-56 | Low |
| Csn1s2a | 1.371 | 1.0E-35 | Low |
| Mylk | 1.243 | 5.5E-43 | Low |
| Wfdc18 | 1.205 | 8.7E-18 | Low |
| Csn2 | 1.192 | 1.2E-40 | Low |
| Synpo2 | 1.107 | 7.0E-34 | Low |
| Oxtr | 1.106 | 7.9E-27 | Low |
| Zfp36 | 1.096 | 1.2E-31 | Low |
| Tsc22d1 | 1.076 | 1.2E-27 | Low |
| Fosb | 1.066 | 6.8E-39 | Low |
| Dusp1 | 1.066 | 1.8E-31 | Low |
| Synm | 1.056 | 3.8E-33 | Low |
| Btg2 | 1.031 | 2.7E-34 | Low |
| Atf3 | 1.029 | 9.2E-36 | Low |
| Hspa1b | 1.028 | 1.1E-38 | Low |
| Zfp36l1 | 1.019 | 2.0E-38 | Low |
| Csn1s1 | 0.999 | 4.9E-35 | Low |
| Cav1 | 0.997 | 2.1E-29 | Low |
| Myh11 | 0.976 | 3.3E-31 | Low |
| Hspa1a | 0.976 | 4.3E-33 | Low |
| Klf2 | 0.965 | 8.6E-28 | Low |
| Tppp3 | 0.944 | 1.9E-22 | Low |
| Cnn1 | 0.940 | 1.1E-25 | Low |
| Ppp1r14a | 0.937 | 4.7E-35 | Low |
| Adamts1 | 0.929 | 2.3E-29 | Low |
| Rras | 0.926 | 8.2E-41 | Low |
| Gadd45g | 0.923 | 6.1E-29 | Low |
| Tns1 | 0.895 | 3.7E-34 | Low |
| Myl9 | 0.878 | 6.8E-29 | Low |
| Jun | 0.867 | 4.0E-29 | Low |
| Actg2 | 0.865 | 1.2E-24 | Low |
| Acta2 | 0.858 | 3.9E-27 | Low |
| Csn1s2b | 0.857 | 1.6E-29 | Low |
| Ckb | 0.825 | 6.4E-31 | Low |
| Tiparp | 0.819 | 3.9E-26 | Low |
| Bhlhe40 | 0.818 | 3.9E-28 | Low |
| Mtss1 | 0.795 | 6.1E-36 | Low |
| Tpm2 | 0.782 | 1.4E-15 | Low |
| Mme | 0.762 | 4.9E-17 | Low |
| Socs3 | 0.760 | 1.2E-11 | Low |
| Ubc | 0.755 | 8.5E-23 | Low |
| Tagln | 0.749 | 2.9E-21 | Low |

**Table S3.** Top 50 differentially expressed genes between siID4 and Non-targeting control Comma-Dβ cells.

| Down in siID4 | | |
|---|---|---|
| Symbol | logFC | FDR |
| Rpl26 | -0.7936339 | 0.00344429 |
| Cers2 | -0.5951326 | 0.00639117 |
| Btf3 | -0.7701147 | 0.00639117 |
| Gclm | -0.7345774 | 0.00639117 |
| Acsl4 | -0.5969801 | 0.00639117 |
| Slc16a1 | -1.0791865 | 0.00639117 |
| Gm15772 | -0.7923647 | 0.00639117 |
| Cd151 | -0.7439322 | 0.00660067 |
| Ywhah | -1.0592157 | 0.00694435 |
| Rap2a | -0.6912214 | 0.00694435 |
| Tpi1 | -0.2935988 | 0.00842821 |
| Mib1 | -0.3095738 | 0.00966817 |
| Arpc1b | -0.5018399 | 0.00966817 |
| Vps37a | -0.7532575 | 0.00966817 |
| Itgav | -0.5716437 | 0.00970215 |
| Ntn1 | -0.7626755 | 0.00995633 |
| Morf4l2 | -0.6064665 | 0.00995633 |
| Adcy7 | -0.2715229 | 0.00995633 |
| Sgms1 | -0.4775586 | 0.00995633 |
| B230219D22R | -0.5082982 | 0.00995633 |
| Adgra2 | -0.5478022 | 0.01103555 |
| Oaf | -0.640244 | 0.01103555 |
| AU019823 | -0.5645291 | 0.01103555 |
| Rock2 | -0.319484 | 0.01237446 |
| Igf2r | -0.25408 | 0.01237446 |
| Slc12a2 | -0.5384233 | 0.01237446 |
| Ptprj | -0.3924337 | 0.01237446 |
| Gfpt1 | -0.4912484 | 0.01237446 |
| Kif23 | -0.5297167 | 0.01237446 |
| Uggt1 | -0.552274 | 0.01237446 |
| Aida | -0.3977576 | 0.01237446 |
| Fdps | -0.3359654 | 0.01237446 |
| Atp5l | -0.8674591 | 0.01258238 |
| Sept8 | -0.3300629 | 0.01271719 |
| Pdia6 | -0.475748 | 0.01312893 |
| Ptpn21 | -0.3673582 | 0.01312893 |
| Vat1 | -0.9032374 | 0.01312893 |
| Slc30a7 | -0.3772582 | 0.01312893 |
| Gas2l3 | -0.6497628 | 0.01312893 |
| Gm12435 | -4.868344 | 0.01312893 |
| Epn2 | -0.3877898 | 0.01322556 |
| Abhd16a | -0.5210536 | 0.01322556 |
| Fam92a | -0.3518148 | 0.01322556 |
| Usp12 | -0.4848923 | 0.01322556 |
| Pmvk | -0.4094487 | 0.0134876 |

| Up in siID4 | | |
|---|---|---|
| Symbol | logFC | FDR |
| Igfbp7 | 0.7103708 | 0.00831843 |
| Id3 | 1.5805303 | 0.00966817 |
| Sparc | 0.3082396 | 0.00966817 |
| Sdc3 | 0.5974385 | 0.00966817 |
| Tmem150a | 0.8564786 | 0.00966817 |
| Cacna1g | 1.5311979 | 0.00970215 |
| Gaa | 0.4475291 | 0.0100761 |
| Col1a1 | 0.6386052 | 0.01103555 |
| Uhmk1 | 0.3365338 | 0.01103555 |
| Slc24a3 | 0.7952255 | 0.01103555 |
| Igfbp2 | 1.336841 | 0.011639 |
| Psmb8 | 0.6798721 | 0.01224243 |
| Plekhb1 | 0.654883 | 0.01237446 |
| Ifitm10 | 0.8621382 | 0.01237446 |
| Trabd2b | 0.6202887 | 0.01312893 |
| Tmeff2 | 1.3380467 | 0.01322556 |
| Tmeff1 | 0.5679847 | 0.01322556 |
| Psmb10 | 0.5993998 | 0.01322556 |
| Cdh15 | 1.2715205 | 0.01322556 |
| Jarid2 | 0.5342555 | 0.01322556 |
| Serf1 | 0.5360454 | 0.0134876 |
| Zcwpw1 | 1.0331659 | 0.01401592 |
| Plxnd1 | 0.5109738 | 0.01414197 |
| Lurap1l | 0.3755962 | 0.01414197 |
| Ncam1 | 0.4747454 | 0.01445327 |
| Msrb3 | 0.3714866 | 0.01445327 |
| Enc1 | 0.3320317 | 0.01460501 |
| Tgtp1 | 1.9855733 | 0.01460501 |
| H19 | 0.6787646 | 0.0149736 |
| Gm5454 | 2.3518513 | 0.0149736 |
| Bst2 | 1.2985464 | 0.01506877 |
| Pld3 | 0.6806763 | 0.01539502 |
| Wnt5b | 1.1296646 | 0.01539502 |
| Ddr1 | 0.3738272 | 0.01541103 |
| Gm17052 | 2.5946567 | 0.01541103 |
| Fam195b | 0.3415067 | 0.01544872 |
| Sort1 | 0.369434 | 0.01544872 |
| Wisp1 | 0.8000815 | 0.01547425 |
| Tfap2b | 0.3567958 | 0.01547425 |
| Tpp1 | 0.3734237 | 0.01547425 |
| Kcp | 3.0068445 | 0.01547425 |
| Fgfr4 | 2.1844006 | 0.01553765 |
| Lima1 | 0.2924242 | 0.01553765 |
| Rsph4a | 2.1844006 | 0.01553765 |
| Etfdh | 0.3947072 | 0.01571933 |

| | | |
|---|---|---|
| Fam168b | -0.7561169 | 0.0134876 |
| Gigyf2 | -0.5809359 | 0.0134876 |
| Nfe2l2 | -0.4266921 | 0.01414197 |
| Pdpn | -0.514649 | 0.01414197 |
| H2-DMb2 | -2.4579144 | 0.01414197 |

| | | |
|---|---|---|
| Col5a1 | 0.5700855 | 0.01580801 |
| Nkain2 | 0.7673748 | 0.01593815 |
| Wisp2 | 1.3916775 | 0.01601568 |
| Gjb4 | 0.3707396 | 0.0162449 |
| Oas3 | 1.2217712 | 0.01628185 |

**Table S4.** Proteins identified in ID4 and HEB RIME experiments.

| ID4 RIME | | | | HEB RIME | | | |
|---|---|---|---|---|---|---|---|
| Uniprot | Protein | p-value | Log(FC) | Uniprot | Protein | p-value | Log(FC) |
| P41139 | ID4_MOUSE | 3.84E-15 | 6.486 | Q61286 | HTF4_MOUSE | 0.0005 | 5.194 |
| P27048 | RSMB_MOUSE | 2.29E-06 | 0.823 | Q9R059 | FHL3_MOUSE | 0.0006 | 2.890 |
| Q9D554 | SF3A3_MOUSE | 4.69E-05 | 0.960 | P70288 | HDAC2_MOUSE | 0.0017 | 0.846 |
| P15806 | TFE2_MOUSE | 6.06E-05 | 1.337 | Q9CYH6 | RRS1_MOUSE | 0.0024 | 2.595 |
| Q9R0E1 | PLOD3_MOUSE | 1.89E-04 | 2.264 | Q8CGB6 | TENC1_MOUSE | 0.0048 | 2.912 |
| Q62203 | SF3A2_MOUSE | 2.27E-04 | 2.305 | Q91Z49 | UIF_MOUSE | 0.0058 | 3.179 |
| P62315 | SMD1_MOUSE | 3.36E-04 | 0.578 | P17710 | HXK1_MOUSE | 0.0081 | 1.732 |
| Q8BFW7 | LPP_MOUSE | 5.04E-04 | 1.543 | Q9D868 | PPIH_MOUSE | 0.0081 | 0.633 |
| Q02788 | CO6A2_MOUSE | 5.24E-04 | 0.785 | Q91VR5 | DDX1_MOUSE | 0.0083 | 0.612 |
| Q8C8U0 | LIPB1_MOUSE | 7.14E-04 | 1.491 | Q8VE37 | RCC1_MOUSE | 0.0085 | 1.487 |
| O08810 | U5S1_MOUSE | 7.57E-04 | 0.577 | O70374 | MTG8R_MOUSE | 0.0088 | 1.125 |
| Q9Z1Y4 | TRIP6_MOUSE | 7.85E-04 | 1.430 | P41139 | ID4_MOUSE | 0.0089 | 0.823 |
| Q91WJ8 | FUBP1_MOUSE | 7.89E-04 | 1.067 | Q9CXY6 | ILF2_MOUSE | 0.0094 | 0.801 |
| Q99NB9 | SF3B1_MOUSE | 1.24E-03 | 0.758 | P13864 | DNMT1_MOUSE | 0.0098 | 2.789 |
| P49718 | MCM5_MOUSE | 1.99E-03 | 1.138 | Q3UQ28 | PXDN_MOUSE | 0.0116 | 3.801 |
| Q9Z1M8 | RED_MOUSE | 2.18E-03 | 1.073 | P63154 | CRNL1_MOUSE | 0.0117 | 0.851 |
| P62320 | SMD3_MOUSE | 2.52E-03 | 0.705 | P70460 | VASP_MOUSE | 0.0135 | 2.965 |
| Q61286 | HTF4_MOUSE | 2.60E-03 | 0.896 | Q8K1R3 | PNPT1_MOUSE | 0.0136 | 2.800 |
| P57784 | RU2A_MOUSE | 3.31E-03 | 0.734 | P61957 | SUMO2_MOUSE | 0.0137 | 1.038 |
| Q3U0V1 | FUBP2_MOUSE | 3.55E-03 | 0.959 | Q60973 | RBBP7_MOUSE | 0.0145 | 1.211 |
| Q6P8I4 | PCNP_MOUSE | 3.95E-03 | 1.652 | Q922K7 | NOP2_MOUSE | 0.0174 | 0.478 |
| O88569 | ROA2_MOUSE | 5.12E-03 | 0.572 | Q8CC35 | SYNPO_MOUSE | 0.0186 | 1.514 |
| Q569Z6 | TR150_MOUSE | 5.93E-03 | 0.943 | P49718 | MCM5_MOUSE | 0.0200 | 0.926 |
| Q3TWW8 | SRSF6_MOUSE | 6.19E-03 | 0.785 | A2AR02 | PPIG_MOUSE | 0.0221 | 0.520 |
| Q8CGF7 | TCRG1_MOUSE | 8.18E-03 | 1.055 | Q99K48 | NONO_MOUSE | 0.0227 | 0.616 |
| Q62318 | TIF1B_MOUSE | 9.61E-03 | 0.658 | O08810 | U5S1_MOUSE | 0.0282 | 0.186 |
| Q99020 | ROAA_MOUSE | 1.10E-02 | 0.582 | Q9CRB2 | NHP2_MOUSE | 0.0291 | 0.572 |
| Q6P4T2 | U520_MOUSE | 1.12E-02 | 0.454 | P84089 | ERH_MOUSE | 0.0303 | 2.006 |
| Q6PE01 | SNR40_MOUSE | 1.13E-02 | 0.665 | Q8VHX6 | FLNC_MOUSE | 0.0311 | 0.861 |
| Q6PDQ2 | CHD4_MOUSE | 1.35E-02 | 0.780 | Q8BFW7 | LPP_MOUSE | 0.0326 | 1.929 |
| Q9Z315 | SNUT1_MOUSE | 1.39E-02 | 1.369 | Q9CPP0 | NPM3_MOUSE | 0.0339 | 0.858 |
| Q8BG81 | PDIP3_MOUSE | 1.52E-02 | 1.154 | Q8VDW0 | DX39A_MOUSE | 0.0394 | 0.786 |
| Q8JZX4 | SPF45_MOUSE | 1.75E-02 | 0.842 | Q9CX86 | ROA0_MOUSE | 0.0412 | 0.887 |
| Q6DFW4 | NOP58_MOUSE | 1.78E-02 | 0.756 | Q9CSH3 | RRP44_MOUSE | 0.0420 | 0.939 |
| Q9JKX4 | AATF_MOUSE | 2.36E-02 | 1.103 | Q80Y84 | KDM5B_MOUSE | 0.0432 | 1.523 |
| Q9JMD0 | ZN207_MOUSE | 2.37E-02 | 1.444 | Q91VC3 | IF4A3_MOUSE | 0.0464 | 1.241 |
| P62317 | SMD2_MOUSE | 2.48E-02 | 0.684 | Q91WN1 | DNJC9_MOUSE | 0.0468 | 0.695 |
| Q61739 | ITA6_MOUSE | 3.07E-02 | 0.594 | O09167 | RL21_MOUSE | 0.0485 | 0.369 |
| Q61789 | LAMA3_MOUSE | 3.19E-02 | 1.806 | | | | |
| Q8R081 | HNRPL_MOUSE | 3.26E-02 | 0.721 | | | | |
| Q9CXY6 | ILF2_MOUSE | 3.66E-02 | 0.248 | | | | |
| P13864 | DNMT1_MOUSE | 3.85E-02 | 0.361 | | | | |
| Q99MJ9 | DDX50_MOUSE | 4.03E-02 | 1.143 | | | | |
| P63154 | CRNL1_MOUSE | 4.17E-02 | 1.396 | | | | |
| Q99KP6 | PRP19_MOUSE | 4.41E-02 | 0.613 | | | | |
| Q8BL97 | SRSF7_MOUSE | 4.48E-02 | 0.471 | | | | |
| Q9QZQ8 | H2AY_MOUSE | 4.63E-02 | 1.121 | | | | |
| Q8BTI8 | SRRM2_MOUSE | 4.92E-02 | 0.639 | | | | |

**Table S5.** Genomic regions of Differentially bound HEB ChIP-Seq peaks between siID4 and NT control Comma-Dβ cells.

| Chr | Start | End | Direction | FC | p-value | FDR | Gene (distance to TSS) |
|---|---|---|---|---|---|---|---|
| chr11 | 113603409 | 113603904 | Up | 3.73 | 4.5E-06 | 0.008 | Slc39a11 (-37898) |
| chr15 | 31873192 | 31873479 | Up | 4.96 | 1.4E-05 | 0.012 | Tas2r119 (-303953) |
| chr15 | 52345685 | 52346124 | Up | 4.53 | 3.6E-05 | 0.021 | Slc30a8 (+50352) |
| chr5 | 148323315 | 148323634 | Up | 4.72 | 5.7E-05 | 0.022 | Slc7a1 (+76429) |
| chr1 | 156874863 | 156875246 | Up | 3.64 | 7.6E-05 | 0.022 | Angptl1 (+36493) |
| chr11 | 117646943 | 117647180 | Up | 3.75 | 1.7E-04 | 0.033 | Tnrc6c (-7751) |
| chr2 | 17508626 | 17508842 | Up | 4.69 | 1.9E-04 | 0.033 | Nebl (+222309) |
| chr11 | 119595488 | 119595972 | Up | 3.55 | 2.0E-04 | 0.033 | Nptx1 (-47977) |
| chr1 | 43656299 | 43656587 | Up | 3.91 | 2.1E-04 | 0.033 | 1500015O10Rik (-74159) |
| chr1 | 12547699 | 12548099 | Up | 4.59 | 2.2E-04 | 0.033 | Sulf1 (-144378) |
| chr15 | 72381629 | 72381874 | Up | 4.08 | 2.5E-04 | 0.033 | Col22a1 (-347525) |
| chr3 | 36505978 | 36506185 | Up | 4.09 | 3.0E-04 | 0.037 | Exosc9 (-46524) |
| chr5 | 127421275 | 127421521 | Up | 3.8 | 3.5E-04 | 0.04 | Tmem132c (+179590) |
| chr6 | 133782477 | 133782683 | Up | 4.3 | 6.3E-04 | 0.067 | Kap (+71182) |
| chr5 | 146659175 | 146659410 | Up | 3.9 | 6.8E-04 | 0.067 | Gpr12 (-74054) |
| chrX | 136261684 | 136261941 | Up | 3.05 | 7.2E-04 | 0.067 | Bex3 (-8440) |
| chr1 | 6037617 | 6037830 | Up | 4.31 | 7.7E-04 | 0.067 | Rb1cc1 (-176921) |
| chr11 | 118717385 | 118718136 | Up | 2.61 | 8.1E-04 | 0.067 | Rbfox3 (+191811) |
| chr6 | 19258755 | 19259056 | Up | 3.66 | 8.2E-04 | 0.067 | Ankrd7 (+392556) |
| chr11 | 119072156 | 119072679 | Up | 2.61 | 1.0E-03 | 0.069 | Cbx8 (-31449) |
| chr15 | 47601402 | 47601633 | Up | 3.29 | 1.0E-03 | 0.069 | NONE |
| chr1 | 19571796 | 19572017 | Up | 4.57 | 1.1E-03 | 0.069 | Tfap2b (+362993) |
| chr8 | 127015103 | 127015345 | Up | 3.51 | 1.2E-03 | 0.069 | Pard3 (-48832) |
| chr11 | 121704506 | 121704827 | Up | 4.13 | 1.2E-03 | 0.069 | Ptchd3 (-125580) |
| chr5 | 112134108 | 112134407 | Up | 3.85 | 1.2E-03 | 0.069 | Cryba4 (+118260) |
| chr6 | 97892878 | 97893163 | Up | 3.42 | 1.4E-03 | 0.069 | Mitf (+85963) |
| chr11 | 120739826 | 120740609 | Up | 1.89 | 1.4E-03 | 0.069 | Hmga1-rs1 (-22576) |
| chr5 | 50279514 | 50279782 | Up | 3.53 | 1.4E-03 | 0.069 | Adgra3 (-220642) |
| chr15 | 49212909 | 49213143 | Up | 4.08 | 1.4E-03 | 0.069 | Csmd3 (-421037) |
| chr5 | 138956388 | 138956610 | Up | 3.71 | 1.4E-03 | 0.069 | Pdgfa (+38538) |
| chr1 | 128871013 | 128871237 | Up | 3.56 | 1.4E-03 | 0.069 | Thsd7b (-402219) |
| chr7 | 126203501 | 126203718 | Up | 3.31 | 1.5E-03 | 0.072 | Xpo6 (-3109) |
| chr15 | 58643530 | 58643741 | Up | 3.33 | 1.6E-03 | 0.072 | Fer1l6 (+133588) |
| chrUn_G | 7649 | 7971 | Up | 2.76 | 1.7E-03 | 0.074 | NONE |
| chr4 | 146733209 | 146733519 | Up | 4.34 | 1.7E-03 | 0.074 | Zfp993 (+122403) |
| chr5 | 18334943 | 18335206 | Up | 3.14 | 1.8E-03 | 0.074 | Gnai1 (+25280) |
| chr1 | 5992485 | 5992734 | Up | 4.18 | 1.8E-03 | 0.074 | Rb1cc1 (-222035) |
| chr5 | 25567753 | 25567985 | Up | 3.32 | 1.8E-03 | 0.074 | Cct8l1 (+51802) |
| chr15 | 53359731 | 53359981 | Up | 3.11 | 1.9E-03 | 0.074 | Gm7489 (-525050) |
| chr1 | 43596044 | 43596292 | Up | 4.26 | 1.9E-03 | 0.075 | 1500015O10Rik (-134434) |
| chr2 | 133666256 | 133666600 | Up | 2.86 | 2.0E-03 | 0.075 | Bmp2 (+114269) |
| chr6 | 86120201 | 86120491 | Up | 2.76 | 2.0E-03 | 0.075 | Tgfa (-74877) |
| chr11 | 115520591 | 115520825 | Up | 3.56 | 2.1E-03 | 0.075 | Jpt1 (-6321) |
| chr5 | 74908540 | 74908781 | Up | 4.03 | 2.2E-03 | 0.08 | Lnx1 (-222874) |

| | | | | | | | |
|---|---|---|---|---|---|---|---|
| chr5 | 58413940 | 58414153 | Up | 3.8 | 2.3E-03 | 0.082 | Pcdh7 (+696080) |
| chr15 | 31420813 | 31421075 | Up | 4.1 | 2.5E-03 | 0.085 | Ankrd33b (-53218) |
| chr11 | 117577237 | 117577684 | Up | 2.05 | 2.5E-03 | 0.085 | Tnrc6c (-77352) |
| chr2 | 98331875 | 98332227 | Up | 3.17 | 2.6E-03 | 0.085 | Gm10801 (-330186) |
| chr18 | 62278421 | 62278663 | Up | 3.64 | 2.7E-03 | 0.088 | Htr4 (-45662) |
| chr15 | 73557550 | 73557776 | Up | 3.6 | 2.9E-03 | 0.088 | Dennd3 (+45103) |
| chr11 | 120846902 | 120847292 | Up | 2.58 | 2.9E-03 | 0.088 | Fasn (-22550) |
| chr15 | 33374091 | 33374388 | Up | 2.94 | 2.9E-03 | 0.088 | Cpq (+291111) |
| chr11 | 114297279 | 114297533 | Up | 3 | 3.0E-03 | 0.089 | Rpl38 (-371203) |
| chr5 | 147695504 | 147695784 | Up | 3.21 | 3.0E-03 | 0.089 | Flt1 (+30367) |
| chr11 | 112755133 | 112755599 | Up | 2.36 | 3.1E-03 | 0.089 | Sox9 (-26858) |
| chr11 | 112201798 | 112202021 | Up | 2.95 | 3.2E-03 | 0.089 | Sox9 (-580314) |
| chrX | 31165181 | 31165593 | Up | 2.73 | 3.3E-03 | 0.09 | Gm10487 (-321918) |
| chr11 | 112820552 | 112820763 | Up | 3.01 | 3.4E-03 | 0.09 | Sox9 (+38434) |
| chr15 | 58576112 | 58576380 | Up | 3.32 | 3.4E-03 | 0.09 | Fer1l6 (+66198) |
| chr11 | 121610031 | 121610403 | Up | 2.89 | 3.7E-03 | 0.095 | Zfp750 (-90884) |
| chr5 | 144711751 | 144712167 | Up | 2.49 | 3.8E-03 | 0.095 | Tmem130 (+49690) |
| chr1 | 10448821 | 10449090 | Up | 3.97 | 3.8E-03 | 0.095 | Arfgef1 (-216286) |
| chr11 | 110257395 | 110257628 | Up | 2.84 | 3.9E-03 | 0.096 | Abca6 (-5736) |
| chr4 | 60468535 | 60468908 | Up | 2.93 | 4.1E-03 | 0.1 | Mup9 (-46788) |
| chr5 | 3853709 | 3853947 | Up | 3.85 | 4.2E-03 | 0.1 | Lrrd1 (+8655) |
| chr6 | 111204465 | 111204679 | Up | 3.65 | 4.4E-03 | 0.103 | Grm7 (+558991) |
| chr11 | 109682162 | 109682504 | Up | 2.48 | 4.4E-03 | 0.103 | Prkar1a (+32928) |
| chr11 | 118063879 | 118064262 | Up | 2.6 | 4.5E-03 | 0.104 | Dnah17 (+65148) |
| chr1 | 85344854 | 85345132 | Up | 1.82 | 4.8E-03 | 0.108 | C130026I21Rik (-74450) |
| chr1 | 171542345 | 171542613 | Up | 3.03 | 5.0E-03 | 0.112 | Cd244 (-16714) |
| chr11 | 117374011 | 117374312 | Up | 2.9 | 5.1E-03 | 0.112 | Gm11733 (-110206) |
| chr5 | 54860627 | 54860836 | Up | 3.82 | 5.6E-03 | 0.119 | Stim2 (+862233) |
| chr19 | 42801489 | 42801721 | Up | 2.76 | 5.6E-03 | 0.119 | Hps1 (-21629) |
| chr1 | 42933785 | 42934078 | Up | 2.59 | 5.7E-03 | 0.119 | Gpr45 (-19031) |
| chr15 | 59334829 | 59335138 | Up | 2.67 | 5.8E-03 | 0.119 | Sqle (+19903) |
| chr11 | 113056112 | 113056519 | Up | 2.31 | 5.9E-03 | 0.119 | Sox9 (+274092) |
| chr11 | 118924971 | 118925289 | Up | 2.31 | 5.9E-03 | 0.119 | Enpp7 (-63058) |
| chr11 | 117586618 | 117586900 | Up | 2.78 | 6.1E-03 | 0.122 | Tnrc6c (-68054) |
| chr6 | 120983229 | 120983464 | Up | 3.39 | 6.2E-03 | 0.122 | Bid (-66494) |
| chr7 | 31481132 | 31481427 | Up | 2.46 | 6.4E-03 | 0.126 | Scgb1b3 (+105688) |
| chr4 | 136110633 | 136110935 | Up | 3.06 | 6.8E-03 | 0.132 | Rpl11 (-57391) |
| chr15 | 59263833 | 59264287 | Up | 2.67 | 7.5E-03 | 0.141 | Mtss1 (-182034) |
| chr17 | 3010105 | 3010352 | Up | 3.18 | 7.8E-03 | 0.144 | Scaf8 (-104743) |
| chr17 | 44999559 | 44999819 | Up | 2.59 | 7.9E-03 | 0.144 | Runx2 (-184892) |
| chr15 | 94727710 | 94727935 | Up | 2.47 | 8.1E-03 | 0.144 | Tmem117 (+98638) |
| chr11 | 115848993 | 115849261 | Up | 2.61 | 8.1E-03 | 0.144 | Myo15b (-9279) |
| chr6 | 37399072 | 37399335 | Up | 2.67 | 8.2E-03 | 0.144 | Dgki (-99230) |
| chr11 | 114254729 | 114255018 | Up | 2.67 | 8.2E-03 | 0.144 | Rpl38 (-413735) |
| chr11 | 113653902 | 113654224 | Up | 2.17 | 8.6E-03 | 0.149 | Cog1 (+4894) |
| chr4 | 61562378 | 61562670 | Up | 1.78 | 8.8E-03 | 0.15 | Mup16 (-42993) |
| chr6 | 97732986 | 97733222 | Up | 2.45 | 8.9E-03 | 0.151 | Frmd4b (-245299) |
| chr1 | 40915275 | 40915499 | Up | 3.11 | 9.1E-03 | 0.152 | Tmem182 (+109786) |

| | | | | | | | |
|---|---|---|---|---|---|---|---|
| chr11 | 111442751 | 111442993 | Up | 2.48 | 9.6E-03 | 0.156 | Kcnj2 (+376708) |
| chr5 | 28864453 | 28864703 | Up | 2.89 | 9.6E-03 | 0.156 | Shh (-397322) |
| chr11 | 121574132 | 121574440 | Up | 2.75 | 9.6E-03 | 0.156 | Zfp750 (-54953) |
| chr5 | 145533568 | 145533817 | Up | 2.18 | 9.7E-03 | 0.157 | Cyp3a16 (-63970) |
| chr11 | 112763431 | 112763699 | Up | 3.56 | 1.0E-02 | 0.161 | Sox9 (-18659) |
| chr11 | 36418901 | 36419112 | Up | 2.82 | 1.0E-02 | 0.161 | Wwc1 (-438480) |
| chr13 | 68863444 | 68863688 | Up | 2.78 | 1.1E-02 | 0.164 | Adcy2 (+135975) |
| chr11 | 110520124 | 110520410 | Up | 3.3 | 1.1E-02 | 0.164 | Kcnj16 (-447766) |
| chr8 | 20379978 | 20380226 | Up | 3.5 | 1.1E-02 | 0.164 | Defb33 (-512567) |
| chr8 | 68625775 | 68626156 | Up | 1.58 | 1.1E-02 | 0.164 | Csgalnact1 (+109180) |
| chr5 | 15498630 | 15498997 | Up | 2.78 | 1.1E-02 | 0.164 | Gm10354 (-519879) |
| chr14 | 76671521 | 76671835 | Up | 2.5 | 1.1E-02 | 0.164 | Serp2 (-114991) |
| chr1 | 194103285 | 194103510 | Up | 3.24 | 1.1E-02 | 0.164 | Camk1g (-733100) |
| chr4 | 3064268 | 3064480 | Up | 2.86 | 1.1E-02 | 0.164 | Vmn1r2 (-107709) |
| chr6 | 62198999 | 62199220 | Up | 2.74 | 1.1E-02 | 0.164 | NONE |
| chr11 | 114226887 | 114227146 | Up | 2.56 | 1.2E-02 | 0.164 | Rpl38 (-441592) |
| chr11 | 120417823 | 120418136 | Up | 1.95 | 1.2E-02 | 0.164 | Faap100 (-39216) |
| chr5 | 30444604 | 30444817 | Up | 2.85 | 1.2E-02 | 0.164 | Otof (+17221) |
| chr11 | 111258367 | 111258730 | Up | 2.43 | 1.2E-02 | 0.164 | Kcnj2 (+192385) |
| chr5 | 73782863 | 73783127 | Up | 2.55 | 1.2E-02 | 0.164 | Spata18 (+131616) |
| chr5 | 20300434 | 20300675 | Up | 2.48 | 1.2E-02 | 0.166 | Phtf2 (+581569) |
| chr11 | 114644379 | 114644645 | Up | 2.69 | 1.2E-02 | 0.166 | Sdk2 (-578294) |
| chr11 | 102067627 | 102067859 | Up | 2.24 | 1.2E-02 | 0.166 | Mpp2 (+20772) |
| chr15 | 61147090 | 61147324 | Up | 2.08 | 1.2E-02 | 0.166 | Myc (-838184) |
| chr6 | 85529458 | 85529741 | Up | 2.44 | 1.2E-02 | 0.166 | Alms1 (-57931) |
| chr1 | 4176403 | 4176771 | Up | 2.51 | 1.3E-02 | 0.168 | Xkr4 (-505089) |
| chr7 | 102953138 | 102953548 | Up | 2.13 | 1.3E-02 | 0.168 | Olfr575 (+2277) |
| chr11 | 120026667 | 120026936 | Up | 2.58 | 1.4E-02 | 0.176 | Aatk (+20268) |
| chr6 | 133131632 | 133131882 | Up | 2.47 | 1.4E-02 | 0.176 | 5530400C23Rik (-160459) |
| chr6 | 90104330 | 90104547 | Up | 2.54 | 1.4E-02 | 0.176 | Vmn1r51 (-18204) |
| chr11 | 113087166 | 113087540 | Up | 1.87 | 1.4E-02 | 0.176 | Sox9 (+305129) |
| chr1 | 22854519 | 22854733 | Up | 2.27 | 1.4E-02 | 0.176 | Rims1 (-48902) |
| chr6 | 135597346 | 135597714 | Up | 2.33 | 1.5E-02 | 0.18 | Emp1 (+234985) |
| chr17 | 86879807 | 86880080 | Up | 2.52 | 1.5E-02 | 0.18 | Tmem247 (-37404) |
| chr11 | 118812535 | 118812791 | Up | 2.29 | 1.5E-02 | 0.182 | Rbfox3 (+96909) |
| chr11 | 116382299 | 116382523 | Up | 2.42 | 1.6E-02 | 0.19 | Foxj1 (-47012) |
| chr1 | 6669297 | 6669577 | Up | 2.25 | 1.6E-02 | 0.19 | Pcmtd1 (-419483) |
| chr1 | 44185102 | 44185333 | Up | 2.14 | 1.6E-02 | 0.193 | Mettl21e (+33743) |
| chr11 | 3132160 | 3132423 | Up | 2.54 | 1.7E-02 | 0.193 | Sfi1 (+61115) |
| chr11 | 120248133 | 120248753 | Up | 1.92 | 1.7E-02 | 0.193 | Bahcc1 (+15496) |
| chr9 | 114303132 | 114303460 | Up | 2.47 | 1.7E-02 | 0.193 | Bcl2a1c (-26839) |
| chr15 | 36189181 | 36189449 | Up | 2.54 | 1.7E-02 | 0.193 | Spag1 (+9947) |
| chr11 | 117633524 | 117634348 | Up | 1.76 | 1.7E-02 | 0.193 | Tnrc6c (-20877) |
| chr1 | 9072878 | 9073087 | Up | 2.26 | 1.7E-02 | 0.193 | Sntg1 (+226255) |
| chr11 | 115895498 | 115895724 | Up | 3.02 | 1.7E-02 | 0.194 | Smim5 (-4355) |
| chr5 | 100575736 | 100576001 | Up | 2.05 | 1.7E-02 | 0.194 | Plac8 (-12446) |
| chr1 | 38176270 | 38176525 | Up | 2.31 | 1.8E-02 | 0.195 | Rev1 (-46736) |
| chr2 | 33703531 | 33703742 | Up | 2.5 | 1.8E-02 | 0.195 | Lmx1b (-63126) |

| | | | | | | | |
|---|---|---|---|---|---|---|---|
| chr11 | 110580970 | 110581207 | Up | 2.91 | 1.8E-02 | 0.196 | Kcnj16 (-386944) |
| chr6 | 112032537 | 112032799 | Up | 1.95 | 1.8E-02 | 0.198 | Lmcd1 (-241090) |
| chr1 | 92265086 | 92265310 | Up | 3.53 | 1.9E-02 | 0.199 | Hdac4 (-84857) |
| chr6 | 119746538 | 119746770 | Up | 1.92 | 1.9E-02 | 0.199 | Wnt5b (-202307) |
| chr11 | 116227525 | 116227809 | Up | 2.37 | 1.9E-02 | 0.199 | Acox1 (-28622) |
| chr5 | 139002011 | 139002401 | Up | 2.06 | 1.9E-02 | 0.2 | Pdgfa (-7169) |
| chr17 | 57748706 | 57748915 | Up | 2.32 | 1.9E-02 | 0.2 | Vmn2r120 (-203497) |
| chr11 | 119536982 | 119537265 | Up | 2.42 | 1.9E-02 | 0.2 | Nptx1 (+10629) |
| chr15 | 59054905 | 59055235 | Up | 2.23 | 2.0E-02 | 0.202 | Mtss1 (+26956) |
| chr1 | 85246334 | 85246580 | Up | 2.02 | 2.0E-02 | 0.205 | A530032D15Rik (-136604) |
| chr1 | 45760764 | 45761110 | Up | 2.52 | 2.0E-02 | 0.205 | Col5a2 (-257655) |
| chr11 | 112667826 | 112668131 | Up | 2.35 | 2.0E-02 | 0.205 | Sox9 (-114245) |
| chr11 | 110076847 | 110077071 | Up | 2.08 | 2.1E-02 | 0.209 | Abca8b (-81114) |
| chr17 | 11093101 | 11093323 | Up | 2.18 | 2.1E-02 | 0.209 | Park2 (+252828) |
| chr15 | 39645924 | 39646131 | Up | 2.42 | 2.1E-02 | 0.209 | Dcstamp (-99904) |
| chr5 | 128753762 | 128754183 | Up | 1.76 | 2.3E-02 | 0.223 | Piwil1 (+17799) |
| chr11 | 120446178 | 120446479 | Up | 1.43 | 2.3E-02 | 0.225 | Tspan10 (+3685) |
| chr2 | 153458118 | 153458369 | Up | 2.37 | 2.3E-02 | 0.226 | 4930404H24Rik (-34546) |
| chr2 | 103687364 | 103687624 | Up | 2.09 | 2.3E-02 | 0.227 | Nat10 (+73776) |
| chr11 | 120299327 | 120299574 | Up | 1.83 | 2.4E-02 | 0.227 | Actg1 (+49091) |
| chr11 | 100563439 | 100563677 | Up | 3.15 | 2.4E-02 | 0.227 | Cnp (-11346) |
| chr15 | 37555056 | 37555277 | Up | 2.76 | 2.4E-02 | 0.228 | 4930447A16Rik (+129613) |
| chr6 | 4126616 | 4126844 | Up | 2.74 | 2.4E-02 | 0.228 | Col1a2 (-378763) |
| chr5 | 67105916 | 67106218 | Up | 2.05 | 2.4E-02 | 0.228 | Tmem33 (-154498) |
| chr11 | 110144486 | 110144894 | Up | 2.1 | 2.5E-02 | 0.228 | Abca8a (-48753) |
| chr15 | 67110611 | 67110986 | Up | 1.96 | 2.5E-02 | 0.228 | Ndrg1 (-141159) |
| chr15 | 83795647 | 83795906 | Up | 2.1 | 2.5E-02 | 0.228 | Mpped1 (+10948) |
| chr11 | 118207409 | 118207748 | Up | 2.33 | 2.5E-02 | 0.228 | Dnah17 (-78360) |
| chr11 | 118321882 | 118322091 | Up | 2.02 | 2.5E-02 | 0.228 | Usp36 (-31756) |
| chr13 | 93728207 | 93728430 | Up | 2.1 | 2.5E-02 | 0.228 | Arsb (-43311) |
| chr11 | 112288324 | 112288546 | Up | 2.19 | 2.5E-02 | 0.228 | Sox9 (-493789) |
| chrX | 94970527 | 94970826 | Up | 2.1 | 2.6E-02 | 0.229 | Spin4 (+56005) |
| chr5 | 73259919 | 73260265 | Up | 1.98 | 2.6E-02 | 0.23 | Fryl (-3473) |
| chr11 | 109982037 | 109982261 | Up | 2.55 | 2.6E-02 | 0.23 | 1700012B07Rik (-154103) |
| chr5 | 37287956 | 37288182 | Up | 2.84 | 2.6E-02 | 0.23 | Crmp1 (+45989) |
| chr1 | 13836247 | 13836552 | Up | 2.06 | 2.6E-02 | 0.23 | Xkr9 (+167629) |
| chr11 | 118690095 | 118690432 | Up | 2.25 | 2.6E-02 | 0.23 | Engase (+213435) |
| chr11 | 114300270 | 114300488 | Up | 2.28 | 2.6E-02 | 0.23 | Rpl38 (-368230) |
| chr11 | 118919826 | 118920315 | Up | 1.75 | 2.7E-02 | 0.231 | Enpp7 (-68117) |
| chr5 | 79440951 | 79441161 | Up | 1.79 | 2.7E-02 | 0.231 | NONE |
| chr8 | 93631226 | 93631526 | Up | 2.33 | 2.7E-02 | 0.231 | Gnao1 (-179937) |
| chr11 | 113527535 | 113527787 | Up | 2.48 | 2.7E-02 | 0.231 | Slc39a11 (+38098) |
| chr11 | 120862627 | 120862849 | Up | 1.91 | 2.7E-02 | 0.231 | Fasn (-38191) |
| chr15 | 35202650 | 35202873 | Up | 2.19 | 2.8E-02 | 0.233 | Osr2 (-93336) |
| chr17 | 14610733 | 14610961 | Up | 3.5 | 2.8E-02 | 0.235 | Thbs2 (+83415) |
| chr8 | 20630952 | 20631164 | Up | 3.05 | 2.8E-02 | 0.237 | Gm15319 (-267848) |
| chr1 | 36580157 | 36580388 | Up | 2.24 | 2.9E-02 | 0.24 | Sema4c (-22747) |
| chr11 | 115232823 | 115233220 | Up | 1.66 | 2.9E-02 | 0.244 | Grin2c (+34221) |

| | | | | | | | |
|---|---|---|---|---|---|---|---|
| chr13 | 65660022 | 65660457 | Up | 1.58 | 3.0E-02 | 0.244 | Gm10324 (-459307) |
| chr5 | 14933551 | 14933994 | Up | 2.65 | 3.0E-02 | 0.244 | Gm9758 (-18874) |
| chr5 | 139568301 | 139568547 | Up | 2.15 | 3.0E-02 | 0.244 | Uncx (+24930) |
| chr15 | 8321463 | 8321725 | Up | 2.73 | 3.0E-02 | 0.244 | Nipbl (+122869) |
| chr5 | 131132457 | 131132673 | Up | 2.59 | 3.0E-02 | 0.244 | A330070K13Rik (-747934) |
| chr15 | 63340242 | 63340733 | Up | 1.87 | 3.0E-02 | 0.244 | Gsdmc (+468271) |
| chr7 | 43194567 | 43194830 | Up | 2.21 | 3.1E-02 | 0.246 | Zfp936 (+17111) |
| chr5 | 20440063 | 20440359 | Up | 2.27 | 3.1E-02 | 0.246 | Phtf2 (+441913) |
| chr6 | 100553519 | 100553748 | Up | 2.26 | 3.1E-02 | 0.246 | Rybp (-266149) |
| chr11 | 118109129 | 118109390 | Up | 2.29 | 3.1E-02 | 0.246 | Dnah17 (+19959) |
| chr5 | 72656656 | 72656978 | Up | 2.08 | 3.1E-02 | 0.246 | Nipal1 (+9022) |
| chr15 | 32635955 | 32636215 | Up | 2.54 | 3.2E-02 | 0.251 | Sdc2 (-284638) |
| chr1 | 11155397 | 11155607 | Up | 2.35 | 3.2E-02 | 0.251 | A830018L16Rik (-259007) |
| chr9 | 84258555 | 84258795 | Up | 1.99 | 3.3E-02 | 0.253 | Bckdhb (+333530) |
| chr8 | 20499680 | 20499890 | Up | 3.25 | 3.3E-02 | 0.256 | Defb33 (-392884) |
| chr11 | 120962386 | 120962655 | Up | 2.06 | 3.4E-02 | 0.26 | Slc16a3 (+13440) |
| chr11 | 119369591 | 119369830 | Up | 1.88 | 3.4E-02 | 0.261 | Rnf213 (-23389) |
| chr5 | 82277664 | 82277917 | Up | 2.03 | 3.4E-02 | 0.261 | Adgrl3 (+967766) |
| chr11 | 120472237 | 120472722 | Up | 1.84 | 3.5E-02 | 0.264 | Arl16 (-4880) |
| chr11 | 117026354 | 117026572 | Up | 2.34 | 3.5E-02 | 0.264 | Sec14l1 (-88709) |
| chr11 | 118413289 | 118413623 | Up | 2.08 | 3.5E-02 | 0.264 | Lgals3bp (-11364) |
| chr11 | 121170523 | 121170744 | Up | 2.32 | 3.6E-02 | 0.264 | Hexdc (-33801) |
| chr11 | 111882991 | 111883241 | Up | 1.68 | 3.6E-02 | 0.269 | Sox9 (-899108) |
| chr7 | 8358091 | 8358562 | Up | 1.75 | 3.7E-02 | 0.27 | Vmn2r43 (-97728) |
| chr15 | 34692589 | 34692924 | Up | 1.72 | 3.7E-02 | 0.27 | Kcns2 (-144624) |
| chr11 | 103707132 | 103707454 | Up | 2.02 | 3.7E-02 | 0.27 | Gosr2 (-9395) |
| chr5 | 26316896 | 26317162 | Up | 2.32 | 3.7E-02 | 0.27 | Dpp6 (-732170) |
| chr11 | 115308171 | 115308542 | Up | 1.65 | 3.8E-02 | 0.273 | Otop2 (+1194) |
| chr5 | 80700937 | 80701157 | Up | 1.8 | 3.9E-02 | 0.274 | Adgrl3 (-608978) |
| chr11 | 111929380 | 111929666 | Up | 2.3 | 3.9E-02 | 0.274 | Sox9 (-852701) |
| chr5 | 35478573 | 35478782 | Up | 2.07 | 3.9E-02 | 0.274 | Cpz (+47020) |
| chr11 | 118858264 | 118858760 | Up | 1.69 | 3.9E-02 | 0.274 | Rbfox3 (+51060) |
| chr5 | 20733906 | 20734133 | Up | 2.44 | 3.9E-02 | 0.275 | Phtf2 (+148104) |
| chr15 | 62355776 | 62356014 | Up | 2.14 | 4.0E-02 | 0.275 | Myc (+370504) |
| chr11 | 120781819 | 120782281 | Up | 1.69 | 4.0E-02 | 0.275 | Gps1 (-2465) |
| chr5 | 125785829 | 125786076 | Up | 2.19 | 4.0E-02 | 0.275 | Tmem132b (+253566) |
| chr5 | 36259095 | 36259337 | Up | 1.96 | 4.0E-02 | 0.277 | Psapl1 (+55195) |
| chr15 | 39952347 | 39952573 | Up | 2.11 | 4.0E-02 | 0.277 | 9330182O14Rik (-189728) |
| chr11 | 121224768 | 121225124 | Up | 1.99 | 4.0E-02 | 0.277 | Ogfod3 (-20235) |
| chr6 | 5273631 | 5273890 | Up | 2.1 | 4.1E-02 | 0.277 | Pon3 (-17475) |
| chr11 | 110070690 | 110070920 | Up | 1.57 | 4.1E-02 | 0.277 | Abca8b (-74960) |
| chr1 | 70171127 | 70171347 | Up | 2.65 | 4.1E-02 | 0.277 | Vwc2l (-554478) |
| chr8 | 20828699 | 20828928 | Up | 2.32 | 4.1E-02 | 0.277 | Gm15319 (-465604) |
| chrUn_JH | 1899 | 2907 | Up | 0.91 | 4.2E-02 | 0.277 | NONE |
| chr15 | 55980575 | 55980799 | Up | 2.16 | 4.2E-02 | 0.277 | Sntb1 (-73738) |
| chr12 | 104983548 | 104983818 | Up | 1.56 | 4.2E-02 | 0.277 | Clmn (-118674) |
| chr11 | 120605770 | 120606073 | Up | 1.66 | 4.2E-02 | 0.279 | Npb (-2555) |
| chr5 | 136823764 | 136824012 | Up | 1.84 | 4.2E-02 | 0.279 | Col26a1 (+59233) |

| | | | | | | | |
|---|---|---|---|---|---|---|---|
| chr17 | 57594755 | 57594989 | Up | 1.72 | 4.3E-02 | 0.28 | Cntnap5c (-174698) |
| chr8 | 92734531 | 92734764 | Up | 2.13 | 4.3E-02 | 0.28 | Mmp2 (-92643) |
| chr8 | 78051333 | 78051573 | Up | 2.04 | 4.3E-02 | 0.28 | Ednra (-327017) |
| chr2 | 153826309 | 153826630 | Up | 1.98 | 4.3E-02 | 0.28 | Sun5 (+44614) |
| chr8 | 84381703 | 84381928 | Up | 1.52 | 4.3E-02 | 0.28 | Ccdc130 (-111436) |
| chr11 | 47474088 | 47474331 | Up | 1.86 | 4.4E-02 | 0.28 | Sgcd (-94688) |
| chr11 | 114532074 | 114532384 | Up | 2.35 | 4.4E-02 | 0.28 | Sdk2 (-466011) |
| chr11 | 120281415 | 120281676 | Up | 2.16 | 4.4E-02 | 0.28 | Bahcc1 (+48599) |
| chr13 | 23622471 | 23623084 | Up | 1 | 4.4E-02 | 0.28 | Hist1h2be (-1654) |
| chr2 | 27802525 | 27802794 | Up | 1.72 | 4.5E-02 | 0.282 | Col5a1 (-83765) |
| chr4 | 139216534 | 139216769 | Up | 2.1 | 4.5E-02 | 0.285 | Capzb (+22656) |
| chr5 | 131065258 | 131065532 | Up | 1.87 | 4.6E-02 | 0.287 | A330070K13Rik (-680764) |
| chr7 | 137708146 | 137708376 | Up | 2.08 | 4.6E-02 | 0.289 | Glrx3 (+270647) |
| chr11 | 110385507 | 110385752 | Up | 1.8 | 4.7E-02 | 0.29 | Abca5 (-47952) |
| chr7 | 43199763 | 43200007 | Up | 1.83 | 4.7E-02 | 0.291 | Zfp936 (+22297) |
| chr1 | 15520675 | 15520932 | Up | 2.14 | 4.8E-02 | 0.294 | Terf1 (-284842) |
| chr1 | 16606818 | 16607118 | Up | 2.18 | 4.8E-02 | 0.294 | Stau2 (-87662) |
| chr4 | 71131358 | 71131720 | Up | 1.44 | 4.9E-02 | 0.294 | Megf9 (-596611) |
| chr11 | 120917505 | 120917997 | Up | 1.95 | 4.9E-02 | 0.294 | Fasn (-93204) |
| chr11 | 111434884 | 111435351 | Up | 1.7 | 4.9E-02 | 0.294 | Kcnj2 (+368954) |
| chr15 | 34849297 | 34849600 | Up | 2.15 | 4.9E-02 | 0.294 | Kcns2 (+12068) |
| chr7 | 130330548 | 130330767 | Up | 1.92 | 4.9E-02 | 0.294 | Fgfr2 (-64467) |
| chr5 | 101419282 | 101419498 | Up | 1.98 | 4.9E-02 | 0.294 | Nkx6-1 (+245606) |
| chr15 | 37386317 | 37386527 | Up | 2.24 | 4.9E-02 | 0.294 | 4930447A16Rik (-39132) |
| chr1 | 184913596 | 184913814 | Up | 2.03 | 4.9E-02 | 0.294 | C130074G19Rik (-30487) |
| chr15 | 27034208 | 27034644 | Up | 2.05 | 4.9E-02 | 0.294 | Ank (-432251) |
| chr11 | 115560890 | 115561366 | Up | 1.71 | 5.0E-02 | 0.295 | Nup85 (-3306) |
| chr14 | 43334403 | 43334700 | Down | -4.08 | 7.2E-05 | 0.022 | Gm8127 (-40739) |
| chr7 | 121419478 | 121419793 | Down | -4.06 | 2.4E-04 | 0.033 | Hs3st2 (+27346) |
| chr11 | 114301903 | 114302223 | Down | -4.03 | 1.0E-03 | 0.069 | Rpl38 (-366546) |
| chr5 | 30536220 | 30536451 | Down | -3.44 | 1.2E-03 | 0.069 | Cib4 (+9500) |
| chr8 | 20827482 | 20827762 | Down | -3 | 3.1E-03 | 0.089 | Gm15319 (-464412) |
| chr6 | 89378435 | 89378652 | Down | -3.02 | 3.2E-03 | 0.09 | Plxna1 (-15924) |
| chr11 | 32357751 | 32357995 | Down | -2.99 | 3.4E-03 | 0.09 | Ubtd2 (-97497) |
| chr15 | 34340243 | 34340513 | Down | -2.64 | 5.5E-03 | 0.119 | 9430069I07Rik (+16043) |
| chr5 | 85845133 | 85845411 | Down | -2.47 | 6.9E-03 | 0.133 | Cenpc1 (+220311) |
| chr5 | 138762198 | 138762543 | Down | -2.62 | 7.2E-03 | 0.136 | Gm5294 (-57709) |
| chr6 | 140132186 | 140132425 | Down | -2.76 | 8.6E-03 | 0.149 | Plekha5 (-291748) |
| chrX | 3399982 | 3400263 | Down | -2.67 | 1.4E-02 | 0.176 | Btbd35f11 (+43567) |
| chr5 | 65172014 | 65172299 | Down | -2.45 | 1.4E-02 | 0.176 | Wdr19 (-27539) |
| chr9 | 106627941 | 106628183 | Down | -2.87 | 1.4E-02 | 0.176 | Iqcf6 (+1480) |
| chr5 | 32291829 | 32292197 | Down | -2.14 | 1.6E-02 | 0.19 | Ppp1cb (-166830) |
| chr15 | 12095958 | 12096175 | Down | -2.5 | 1.8E-02 | 0.195 | Sub1 (-100009) |
| chr5 | 19931121 | 19931378 | Down | -2.42 | 1.8E-02 | 0.199 | Magi2 (+704199) |
| chr11 | 113631918 | 113632226 | Down | -2.12 | 2.1E-02 | 0.209 | Cog1 (-17097) |
| chr8 | 71147476 | 71147740 | Down | -2.38 | 2.5E-02 | 0.228 | Myo9b (-125106) |
| chrX | 75689334 | 75689640 | Down | -1.95 | 2.5E-02 | 0.228 | Rab39b (-111256) |
| chr7 | 14757574 | 14757850 | Down | -2.01 | 3.3E-02 | 0.253 | Obox2 (-631009) |

| | | | | | | | |
|---|---|---|---|---|---|---|---|
| chr5 | 112547684 | 112547928 | Down | -2.11 | 3.5E-02 | 0.264 | Sez6l (+29169) |
| chr11 | 116405070 | 116405373 | Down | -1.59 | 3.7E-02 | 0.27 | Foxj1 (-69823) |
| chr11 | 43447353 | 43447596 | Down | -2.55 | 3.7E-02 | 0.27 | C1qtnf2 (-26831) |
| chr15 | 19515055 | 19515318 | Down | -1.94 | 3.9E-02 | 0.274 | Cdh10 (+694858) |
| chrX | 33988514 | 33988811 | Down | -1.45 | 4.1E-02 | 0.277 | Gm21870 (-1847) |
| chr1 | 6643322 | 6643612 | Down | -2.04 | 4.5E-02 | 0.285 | Pcmtd1 (-445453) |
